# *De novo* design of protein competitors for small molecule immunosensing

**DOI:** 10.64898/2025.12.16.694474

**Authors:** Yosta de Stigter, Tallie Godschalk, Maarten Merkx

## Abstract

Competition-based immunoassays are a common strategy for detecting small-molecule biomarkers. However, these assays rely on the availability of a custom competitor molecule that can effectively be displaced upon analyte binding, often requiring time-consuming synthesis and conjugation steps. *De novo* designed protein binders present a compelling alternative, as their binding properties can be tuned and they allow for straightforward genetic-incorporation into existing immunoassays. Here, we leverage the BindCraft pipeline to design competitive binders by targeting antigen-binding sites, followed by *in silico* filtering to select for steric clashes with the small-molecule analyte. As a proof of concept, we designed digoxin competitors and experimentally screened the binders using a simple bioluminescent assay, identifying 7/10 successful binders directly in bacterial lysate. These binders exhibited low to moderate binding affinities (*K*_*d*_ = 42 nM – 1.1 µM) and were displaced by digoxin. Two *de novo* binders were encoded into a previously established competition-based immunosensor, enabling sensitive digoxin detection (*K*_*d*_ = 109 nM). These results demonstrate that deep learning-based models can rapidly yield effective competitor binders, enabling straightforward adaptation and optimization of sensing platforms for small-molecule targets.

## Introduction

Detection and quantification of small-molecule biomarkers, such as drugs and hormones, is crucial for disease diagnostics, therapeutic monitoring and general health assessment^1–4^. Immunoassays like enzyme-linked immunosorbent assays (ELISA) are widely used for biomarker detection due to their high specificity and sensitivity^5^. However, for small molecules, the absence of multiple antibody binding sites precludes use of the sandwich immunoassay formats used for protein targets. Instead, competitive immunoassay formats are employed, in which the target analyte competes with a labeled analog for antibody binding. Developing immunoassays is not straightforward as it requires extensive optimization of assay conditions and often involves complex, multistep workflows that restricts their use to centralized laboratories, resulting in long sample-to-answer times. In competitive formats specifically, assay performance and adaptability are further constrained by the requirement of a suitable competitor analog.

To address some of these limitations, various homogeneous immunoassays formats have been developed that are more modular and suited for a point-of-care (POC) application^6,7^. One notable example is the small-molecule–responsive antibody switch developed by Soh et al., in which a competitor analog is linked to an analyte-specific antibody through a DNA scaffold that upon displacement causes a change in Förster resonance energy transfer (FRET)^8^. To eliminate the need for external excitation, bioluminescent sensors formats have also been explored, such as the luciferase-based indicators of drugs (LUCIDs) and bioluminescence resonance energy transfer (BRET)-based quench bodies^9–11^. Building on these concepts, our group recently developed a luminescent competition sensor (LUCOS) platform, which includes an universal bioluminescent adapter protein that incorporates a competitor molecule and couples to an antibody through protein G-mediated photoconjugation^12^. Displacement of the competitor by free analyte induces a conformational switch, resulting in a change in bioluminescent emission ratio.

Despite their advantages, these sensor formats remain dependent on the displacement of a competitor molecule, which fundamentally limits their sensitivity and broad applicability^13,14^. Because the competitor typically comprises an analog of the analyte with a similar binding affinity, its displacement is often inefficient. This is especially apparent in intramolecular sensor designs, in which the free analyte must approach the effective concentration of the analog for it to be displaced. Although the affinity of the competitor can be tuned to some extent, for example by changing linker properties or synthesizing structural analogs with weaker binding affinities, this process is analyte-specific and time-consuming^8,11^. Moreover, suitable small-molecule analogs with reactive groups for conjugation are often unavailable and require custom synthesis, further complicating sensor adaptation to new analytes. An universal strategy to generate competitors that can be readily incorporated into sensor designs would therefore highly benefit the development competition-based small-molecule sensors.

Advances in computational protein design are now enabling very efficient and accurate design of protein binders^15–17^. Whereas traditional methods to develop binding domains such as immunization or directed evolution are time-consuming and provide limited control over the targeted epitope, deep learning-based models can rapidly generate protein structures and sequences that are confidently predicted to bind to an user-defined site. A current example of such a method is BindCraft, which leverages the AlphaFold2 structure prediction network for binder design and proteinMPNN for sequence optimization^18,19^.BindCraft demonstrates high experimental success rates across different targets, with most designed binders exhibiting a low to moderate affinity (micromolar- to nanomolar-range). While higher-affinity binding is generally desirable for therapeutic and biotechnological applications, this initial weak binding affinity can be advantageous for application as a competitor molecule.

In this work, we demonstrate the *de novo* design of protein binders to serve as small-molecule competitors (Figure 1). These competitor binders typically display weak binding affinities which enables efficient displacement and allow straightforward incorporation through genetic encoding. Using the BindCraft pipeline, we generate binders targeting the antigen-binding site of small-molecule-specific antibodies, and subsequently filter designs *in silico* based on interaction confidence metrics as well as predicted steric clashes between the designed binder and small molecule analyte to ensure competitive binding. Rapid experimental screening is facilitated by a simple bioluminescent assay based on binding-induced complementation of split luciferase fragments, allowing efficient assessment of binding, competition, and subsequent affinity determination. As a proof of concept, competitor binders were designed against an anti-digoxin antibody. We identified several low- to moderate-affinity competitors and incorporated two into the LUCOS sensor platform to demonstrate digoxin detection. Overall, this work demonstrates how deep learning-based models can rapidly yield effective competitor binders, enabling straightforward adaptation of competition-based small-molecule sensors to different targets.

**Figure 1.**
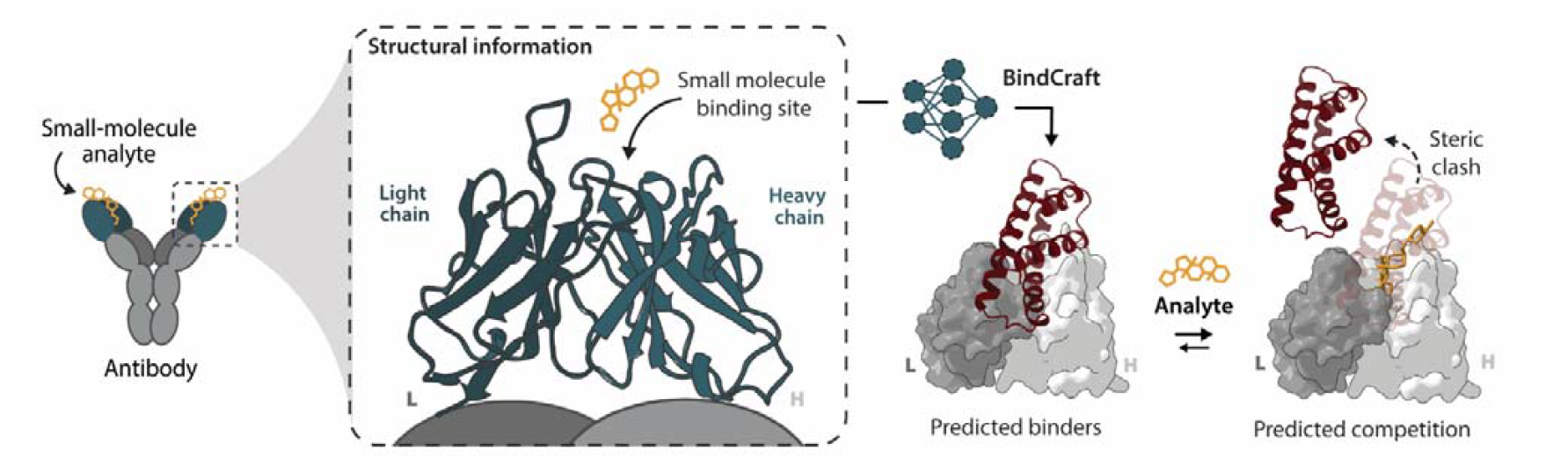
*De novo* design of protein competitors. Using the BindCraft pipeline, competitor binders are designed to target the small-molecule binding site of an antibody. To ensure competitive binding with the small-molecule analyte, binders are selected that are predicted to sterically clash with the analyte.

## Results and discussion

### Competitor design

To design proteins that compete with antibody-small-molecule interactions, we targeted binders towards the antigen-binding site. Antibodies typically bind their small molecule analytes at the interface between the light and heavy chain, with interactions mediated by the structured regions as well as the complementarity-determining regions (CDR)-loops^20^. To promote competitive binding, binders were directed towards interacting residues or neighboring sites that are accessible and preferably hydrophobic as hydrophobic interactions are generally more tractable for computational design^15,21^. These hotspot residues were used as input for the BindCraft pipeline, together with the structure of the target antibody. To reduce computational load, we used an AlphaFold2 (AF2) prediction of a truncated antibody containing just variable domains that are involved in analyte binding. Successful BindCraft designs were filtered using AlphaFold3 (AF3) interaction confidence metrics (iPTM > 0.8), considering that AF3 has been shown to improve antibody docking predictions^22,23^. As overall AF3 interactions metrics can be biased by the high confidence of the light- and heavy-chain contacts, we subsequently employed the AlphaBridge software to independently assess binder-antibody interaction (contact iPTM > 0.75)^24^. Finally, AF3-predicted binder-antibody complexes were overlayed with the corresponding analyte-antibody crystal structures to identify the candidates predicted to sterically clash with the target analyte.

To test this workflow, we selected digoxin as an initial target (Figure 2A). Digoxin is a plant-derived steroid used to treat various cardiac disorders and has a narrow therapeutic window, necessitating accurate monitoring in patients^25^. Anti-digoxin antibodies with known crystal structures are commercially available, eliminating the need for time-consuming antibody development^26^. Because BindCraft relies on AF2 predictions for both design and validation, accurate prediction of the antibody variable regions is critical. AF2 occasionally fails to correctly predict antibody structures, which necessitates known crystal structures to evaluate the prediction quality. For anti-digoxin (pdb: 1IGJ), the AF2 prediction of the variable domain closely matches the crystal structure, confirming its suitability as input target for BindCraft (Figure 2A, RMSD = 0.502 Å). We selected three amino acids involved in digoxin binding as hotspots (light chain: Val99, heavy chain: Tyr32, Trp103) and used them independently as input for separate BindCraft runs, with binder lengths of 80-120 amino acids. Of the 220 initial designs that passed default BindCraft filters, 40 passed the additional filtering with AF3, AlphaBridge and structural overlay. The 10 designs with the highest confidence metrics were selected for experimental validation (supplementary Table S1).

**Figure 2.**
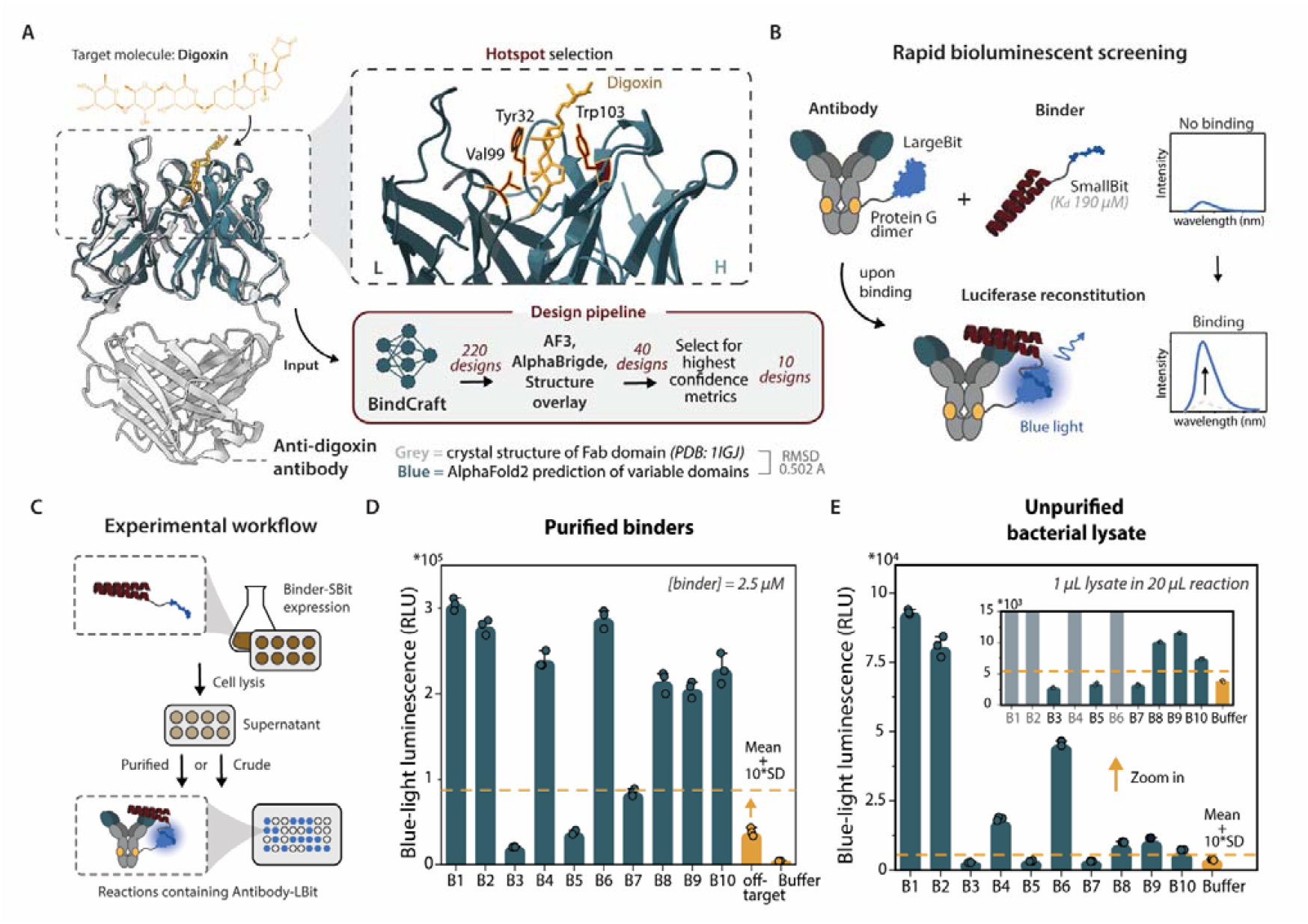
Digoxin competitor design and rapid experimental screening using split luciferase assay. A) Binders are designed to compete with digoxin (orange) for antibody binding. The AF2 prediction of anti-digoxin (26-10, pdb: 1IGJ) variable domain is used as input for BindCraft together with one of three hotspot residues (red). Successful BindCraft designs are filtered with AF3, AlphaBridge and structure overlay to predict steric clashes. B) Bioluminescent screenings assay design. The antibody is fused to NanoLuc LargeBit through a protein G dimer that is photoconjugated to the Fc domain. Binders are fused to the lowest-affinity SmallBit (190 µM). Binding of antibody and binders results in luciferase reconstitution and an increase in blue-light luminescence. C) Workflow of binder screening using both purified binders and clarified lysate as assay input. D,E) Luminescence intensity of the screened binders (B1-B10) using purified binders (D) or bacterial lysate (E). Anti-digoxin-LBit (1 nM) is incubated with 2.5 µM binder (D) or 20x diluted lysate (E) for 30 minutes at room temperature in 1xPBS + 0.1% BSA, followed by the addition of 1000x diluted NanoLuc substrate. Luminescence intensity is measured at 458 nm. Off-target control includes TNFα-SBit (2.5 µM). Dashed lines represents mean + 10* standard deviation of the off-target control (D) or buffer control (E). Inset shows zoomed-in view of the same graph. Bars represent mean luminescence intensity and circles represent individual values with n = 3 technical replicates.

### Bioluminescent binding screenings assays

To enable rapid experimental screening of *de novo* binders, we employed a simple binding assay based on split NanoLuc luciferase complementation (Figure 2B)^27,28^. This luminescent readout does not require external excitation, which typically limits sensitivity in complex matrices due to autofluorescence and scattering, and therefore may allow for direct screening in bacterial lysate, obviating laborious protein purification. To this end, NanoLuc Large Bit (LBit) was covalently coupled to the antibody through a protein-G dimer (pG-LBit) incorporating a photocrosslinkable non-natural amino acid (para-benzoyl-phenylalanine, pBPA), as used for previous bioluminescent immunoassays^29,30^ The designed binders were genetically fused to the lowest-affinity NanoLuc Small Bit (SBit) fragment (*K*_*d*_ = 190 µM) to minimize background complementation for reliable screening of lower-affinity binders. Although SBit and LBit are mutually interchangeable^31^, using the smaller SBiT (1.3 kDa vs 18 kDa for LBiT) as binder fusion partner likely reduces the chance of interference with binding or expression.

The designed binders were encoded into plasmids containing SBit and expressed at small-scale in *Escherichia coli*, followed by chemical lysis and centrifugation to obtain the clarified lysate (Figure 2C). As a positive control, binders were also purified via his-tag affinity chromatography, yielding pure protein in sufficient amounts (∼5-20 mg/mL, supplementary Figure S1). pG-LBit was similarly expressed in *E. coli* but with amber stop codon suppression and tRNA synthetase/tRNA pair co-transformation to incorporate the pBPA amino acid and purified using both his-tag and strep-tag affinity chromatography to obtain pure protein (∼20 mg/mL, supplementary Figure S2). Photocrosslinking was performed by incubating the anti-digoxin antibody and pG-LBit at a 1:2 molar ratio for 30 min and subsequently illuminating the mixture with UV light (30 min, λ = 365 nm). Non-reducing SDS-PAGE analysis suggested successful photoconjugation with minimal unconjugated antibody (supplementary Figure S3). For all proteins, correct protein masses were verified using ESI-Q-ToF mass spectroscopy (supplementary Figure S4 and S5). For the *de novo* binders, circular dichroism spectra showed a characteristic double-dip that indicates that the proteins are well-folded and contain the alpha-helical secondary structure elements consistent with their design (supplementary Figure S6).

To identify successful binders, purified binder-SBit constructs (2.5 µM) were incubated with photoconjugated anti-digoxin antibody-LBit (1 nM) for 30 minutes at room temperature, followed by the addition of NanoLuc substrate. A negative control containing a non-binding protein fused to the same SBit (TNF-α, from earlier work^29^) was included to account for background LBit-SBit complementation at the tested binder concentrations. Binders were considered successful if their luminescence intensity surpassed that of the negative control plus 10 times the standard deviation of the replicates, although this threshold can be adjusted to apply less or more strict filtering. Notably, 7 out of our 10 designed binders displayed luminescence intensities above the set threshold, indicating successful binders (Figure 2D). When using clarified lysate (20x diluted) instead of purified protein as input, the same 7 binders were identified as positive (Figure 2E), demonstrating that protein purification is not strictly necessary for initial screening. Although absolute luminescence intensities were lower in the presence of lysate, binders could still be distinguished from a control without lysate. Several binders displayed a more pronounced increase in luminescence, which reflects differences in binding affinity and/or expression levels among binders. Together, these results shows that the split luciferase assay enables fast screening of functional binders directly from lysate, thereby facilitating the selection of designs for more thorough characterization.

### Characterization of binding and competition

Next, the seven hits were characterized for their binding affinities and competitive binding with digoxin (Figure 3A). Apparent binding affinities were determined using the same split luciferase assay by titrating varying amounts of binders (9.8 nM – 1.25 µM) against a fixed concentration of antibody (1 nM). Reactions were incubated for 30 minutes, followed by the addition of NanoLuc substrate. All binders showed a concentration-dependent increase in luminescence indicative of binding, with apparent affinities ranging from 42 nM to 1.1 µM (Figure 3BC). Please note that the apparent *K*_*d*_ values resulting from the split luciferase assay are approximate and do not reflect true binding affinities as LBit-SBit interactions can influence binding strength depending on the tested binder. To assess whether the binders competitively bind the digoxin binding site, we titrated digoxin (*K*_*d*_ = 9 nM) to the formed binder-Ab complexes. Reactions were incubated for 2 hours prior to luminescence measurement to ensure that equilibrium was reached. All binders showed a decrease in luminescence upon addition of digoxin, indicating successful binder displacement (Figure 3D and supplementary Figure S7). Also, the observed competitive binding suggests that binders interact with the intended epitopes, further confirming the design models (Figure 3E).

**Figure 3.**
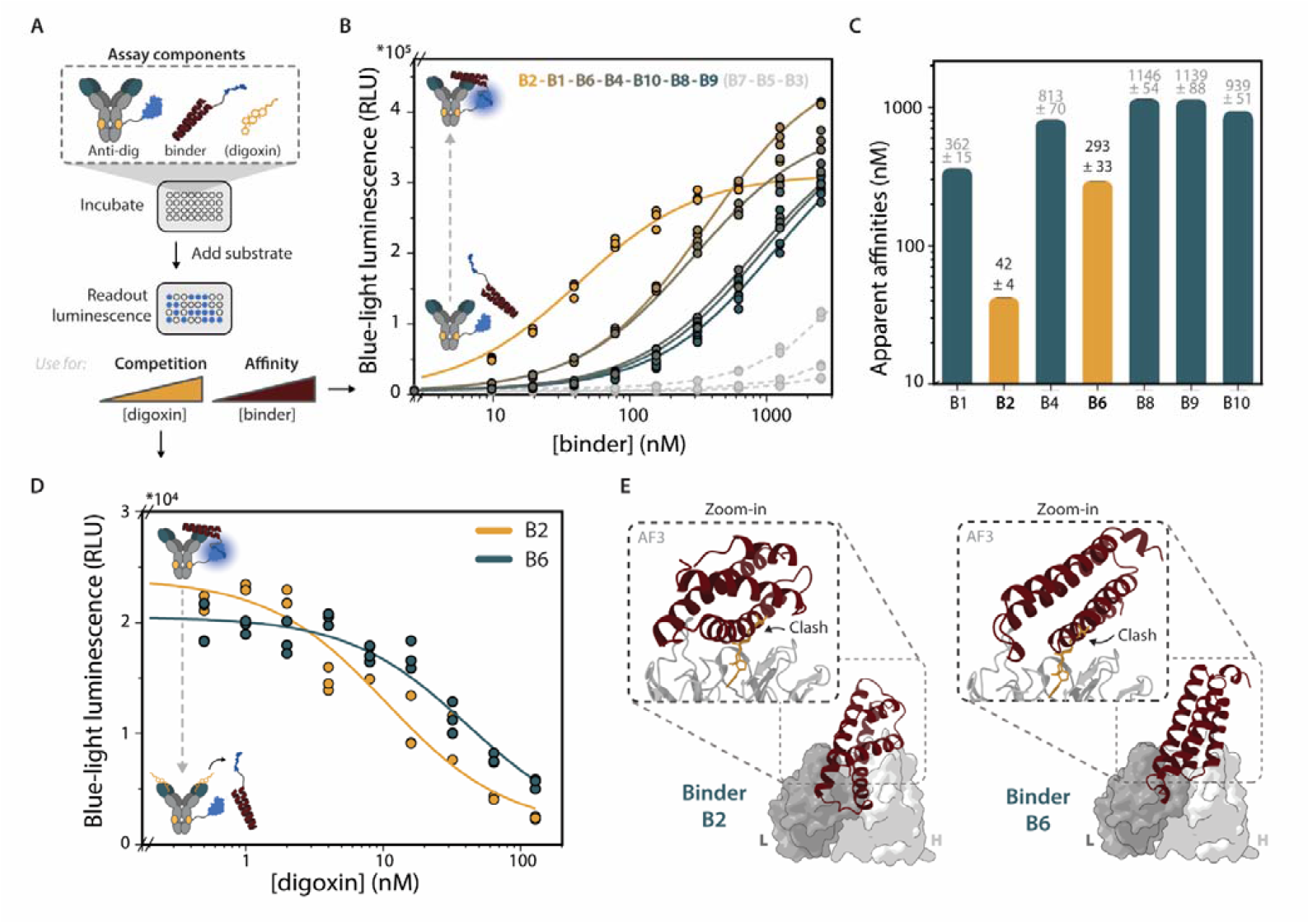
Characterization of binding affinity and competition. A) Schematic overview of split luciferase assay for determining binding affinities and competitive binding. B) Bioluminescence titrations with binder hits to approximate binding affinities. Anti-digoxin-LBit (1 nM) is incubated with 9.8 nM – 1.25 µM binder and incubated for 30 minutes before the addition of NanoLuc substrate. Individual data points (n = 3) are represented as circles and the model fit as solid lines. C) Apparent affinities derived from binding curves displayed in panel B. Bars represent apparent *K*_*d*_ values derived from fitted data (supplementary Figure S8). Apparent *Kd* value ± standard error is indicated above the bars. D) Competitive titrations of binders B2 and B6 with digoxin. Anti-digoxin-LBit (500 pM) was incubated with B2 (5 nM) or B6 (50 nM) for 30 minutes to ensure complex formation. Reactions were then combined with increasing concentrations of digoxin (500 pM – 128 nM) and incubated for 2 hours prior to the addition of NanoLuc substrate. Individual data points (n = 3) are represented as circles and the model fit as solid lines. E) Graphical representation of structures of binders B2 and B6. Drawing of binders (red)-antibody (grey) complexes are based on AF3 predictions. Placement of digoxin (orange) is based on pdb structure 1IGJ.

### LUCOS incorporation

To demonstrate digoxin sensing and evaluate whether *de novo* competitors are compatible with existing sensing platforms, we incorporated the anti-digoxin binders into our previously developed luminescent competition sensor (LUCOS). LUCOS is a modular bioluminescent platform that relies on intramolecular competition between a tethered competitor and free target analyte for antibody binding^12^. Sensors are assembled through site-specific covalent coupling of a bioluminescent adapter protein to a target-specific antibody using protein G-mediated photoconjugation to the Fc domain, as described for the bioluminescent screening assay. This adapter protein consists of protein G, genetically fused to LBit, the competitor domain and SBit (*K*_*d*_ = 2.5 µM). Binding of free target analyte displaces the competitor and promotes formation of the closed conformation of the sensor with reconstituted NanoLuc, resulting in increased blue luminescence. To compensate for signal decay caused by substrate depletion, a green light emitting calibrator luciferase (λ = 530 nm) is used, comprising split NanoLuc genetically fused to mNeonGreen. Analysis of the ratio between blue and green signal enables a time-independent readout and internal sample calibration (Figure 4A)^29^.

**Figure 4.**
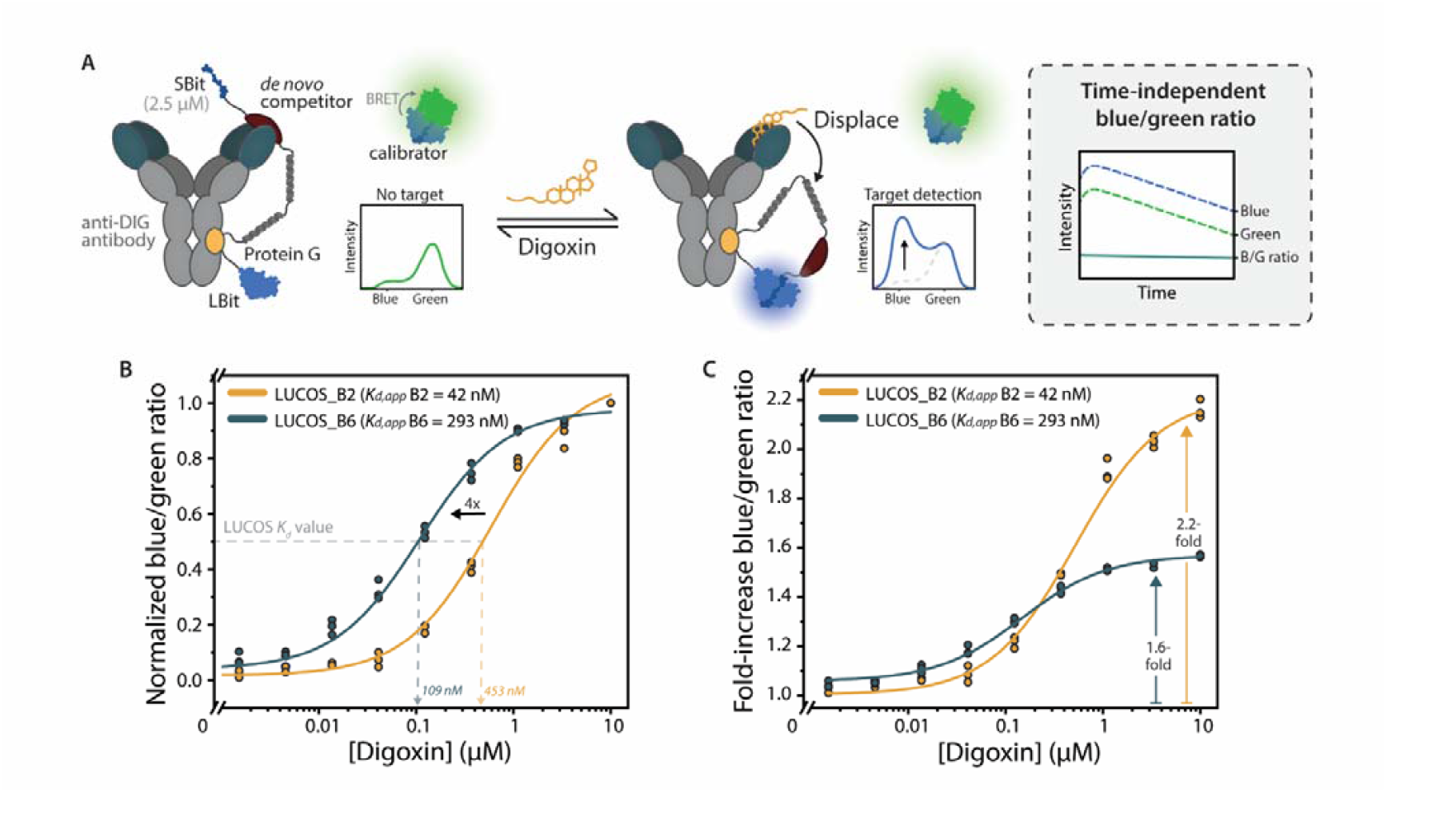
*De novo* designed competitor domains in LUCOS platform. A) LUCOS assay design. The antibody is fused to LUCOS adapter through protein G-mediated photoconjugation. Adapter contains N-terminal NanoLuc LBit and a semi-rigid protein linker connecting to the *de novo* competitor domain, followed by the C-terminal NanoLuc SBit (K_D_ = 2.5 µM). Competitor binding keeps construct in the open state, prohibiting the emission of blue luminescent signal. The split calibrator luciferase produces a constant green luminescent background signal. Target binding induces competitor displacement and closing of the sensor construct, enabling blue signal emission. Taking the ratio between the blue and green signal results in a time-independent constant signal. B) LUCOS titration assay results. LUCOS_B2 or LUCOS_B6 (1 nM) was incubated with digoxin (1.5 nM – 10 µM) for 2 hours to ensure equilibrium prior to addition of NanoLuc substrate. Blue/green ratio was determined by dividing emission at 358 nm by emission at 533 nm. C) Effect of competitor affinity on dynamic range in LUCOS titration assay. Fold-increase in blue/green ratio was determined by dividing the calculated blue/green ratios by the blue/green ratio of the background signal. Individual data points (n = 3) are represented as circles and the model fit as solid lines.

The desired affinity of competitors for LUCOS integration depends on several parameters, including the analyte-antibody affinity and the effective molarities imposed by linker lengths, as well as the required sensitivity range. This involves a delicate balance in which the binders must bind strong enough to bring the sensor into its open state, yet as weakly as possible to ensure effective displacement by the analyte. Thermodynamic modeling suggests that a low-nanomolar (∼ 10 nM) competitor is required to fully shift the sensor into its open state, enabling a digoxin response in the nanomolar to micromolar range (supplementary Figure S9). A weaker competitor (∼ 100 nM) would allow more sensitive detection, albeit with a smaller dynamic range. Guided by these modelling results, we selected the binders B2 and B6 with apparent *K*_*d*_ values of 42 and 293 nM, respectively, to integrate into the LUCOS platform (Figure 3E).

Two LUCOS adapter proteins containing the two respective *de novo* binders were encoded into plasmids and expressed in *E. coli* following the same strategy as described for pG_LBit. Chemical lysis followed by purification of the cleared lysate with Ni^2+^ affinity chromatography followed by Strep-Tactin purification resulted in pure samples with a yield of ∼12 mg/L (B2) and ∼17 mg/L (B6) (supplementary Figure S10). ESI-Q-ToF mass spectrometry confirmed the correct protein masses (supplementary Figure S11). Photoconjugation was performed by incubating the anti-digoxin antibody and adapter proteins in a 1:1 molar ratio for 30 minutes and subsequently illuminating the mixture with UV light for 30 minutes. Non-reducing SDS-PAGE analysis suggested successful coupling of the adapter proteins to the antibody (supplementary Figures S12 and S13).

The performance of the LUCOS sensors was tested in a titration experiment where 1 nM of LUCOS was incubated with 1.5 nM – 10 µM digoxin and 100 pM calibrator luciferase for 1 hour. Integration of both binders resulted in a digoxin-dependent increase in blue/green ratio, suggesting that the binders are effectively displaced by digoxin to enable luciferase reconstitution. The observed response ranges matched the predictions of the thermodynamic model: the apparent *K*_*d*_ of LUCOS for free digoxin using B2 as competitor was 543 ± 90 nM, which improved to an *K*_*d*_ value of 109 ± 11 nM using the lower-affinity competitor B6 (Figure 4B). The increase in sensitivity coincided with a slightly reduced dynamic range, presumably due to incomplete opening of the sensor in the absence of digoxin (Figure 4C and supplementary Figure S14). Nonetheless, these results clearly demonstrate the strength of applying deep learning-based models to design tunable competitor proteins to improve the modularity of competition-based assays for small molecule detection.

## Conclusions

In this work, we established the *de novo* design of competitive binders as an universal strategy for adapting small-molecule sensors to different analytes. Leveraging the BindCraft design pipeline combined with additional computational filtering, *de novo* binders targeting an anti-digoxin antibody were generated with high experimental success rate (70%), eliminating the need for extensive library screening. A split luciferase assay enabled rapid pre-screening of binders from bacterial lysate, followed by affinity determination to select suitable candidates. Successful binders displayed low to moderate binding affinities (42 nM to 1.1 µM) and were displacement by digoxin, rendering them well-suited as competitors for typical high-affinity antibody-analyte pairs. Two digoxin competitors were encoded into the existing LUCOS platform, demonstrating sensitive digoxin detection *(K*_*d*_ = 109 nM) without the need for post-translational conjugation and/or custom competitor synthesis. These results suggest that *de novo* designed binders offer an easy and tunable alternative to traditional analog competitors, enabling straightforward adaptation and optimization of existing assays to different small-molecule targets.

While antibodies are favorable binders due to their high affinity and specificity, their use as target also imposes several limitations. Accurate design of competitors relies on antibody-analyte structures, which are not always available for commercial antibodies, limiting the amount of targets to which the method can currently be applied. Although AF2 has demonstrated an impressive ability to predict protein-protein interactions, antibodies and antibody-antigen complexes remain challenging and can hamper BindCraft use in case of inaccurate predictions. As the scarcity of antibody-antigen structures similarly limits progress in other fields of research, such as in *de novo* antibody design, ongoing efforts to obtain more structural data will also broaden the applicability of our design strategy to a wider range of targets. Furthermore, we expect that improved prediction of small-molecule-antibody interactions, for example using AF3, can help to further automate screening for competition or steric clashes between binder and analyte.

The split luciferase assay enables efficient screening, characterization and selection of binders for downstream applications. The bioluminescent readout allows for measuring binding directly in lysate, eliminating the need for laborious protein purification. Results from lysate and purified proteins were found consistent, yet variation in expression levels and binding affinities can potentially lead to false-positive or - negative results. False positives are effectively mitigated by using diluted lysate and low-affinity SBit. Although false negative results are more difficult to correct for, low expression or weak affinity is generally not desirable and therefore naturally penalized during screening. When interested in lower-affinity interactions, assays using purified protein at defined concentration will provide a more reliable measurement. The split luciferase assay also allows for straightforward approximation of binding affinities. While apparent affinities can be biased by the additional SBiT and LBiT interaction, determining the exact affinities is not strictly necessary for application as a competitor. For other cases, the assay can serve as a rapid pre-screening tool to identify promising binders prior to more laborious characterization techniques such as surface plasmon resonance (SPR) or biolayer interferometry (BLI).

The success of competitor design likely depends on the size and binding orientation of the target analyte. While all digoxin binders showed competitive binding, many clinically-relevant biomarkers such as hormones and drugs are smaller and may therefore be more difficult to displace. Achieving competition in such cases may require more careful design of binders and/or more extensive *in silico* and experimental screening. In future efforts, we aim to showcase the modularity of *de novo* competitor design by applying our method to a wider range of small-molecule targets.

## Methods

### Competitor design and *in silico* filtering

The BindCraft pipeline (https://github.com/martinpacesa/BindCraft) was run on the Snellius HPC, using an H100 GPU. An AF2 prediction of the variable domains of an anti-digoxin antibody (clone 26G10) was used as the target structure input for the BindCraft pipeline (for sequence see supplementary information). Structural accuracy of the target prediction was checked against the full-length analyte-antibody crystal structure (pdb: 1IGJ). Residues involved in digoxin binding were selected as hotspots and used independently for separate BindCraft runs. The BindCraft pipeline was employed to generate binders with a length between 80-120 amino acids, in combination with default filtering settings. AF3 prediction confidence metrics were subsequently used to filter successful BindCraft designs (iPTM > 0.8), followed by the AlphaBridge webtool to assess potential binder-antibody interactions (contact iPTM > 0.75). Competitive binding of designs binders and digoxin was analyzed using overlays with the crystal structure in UCSF ChimeraX.

### Cloning

For screening of the *de novo* designed binders, synthetic, codon-optimized gene fragments encoding the *de novo* binders were ordered from Integrated DNA Technologies (IDT). A previously designed pET28a expression vector encoding a semi-rigid protein linker, SBit peptide (2.5 µM) and C-terminal hexahistidine-tag was linearized and used as backbone for binder insertion through Gibson Assembly using the HiFi DNA assembly kit (New England Biolabs) according to manufacturer’s instructions. The LUCOS adapter proteins containing binders B2 and B6 were cloned via traditional restriction/ligation into the pET28a LUCOS expression vector containing an N-terminal hexahistidine-tag and C-terminal streptavidin-tag. Inserts were obtained through PCR (Phusion, NEB) on the gene fragments, using primers containing overhangs encoding for an N-terminal *Spe*I and C-terminal *Age*I restriction enzyme cut site. Cloned constructs were transformed in *E. coli* TOP10 cells and plated on ager containing 50 µg/mL kanamycin. Small cultures were prepared using 5 mL Lysogeny Broth (LB) medium (10 g/L NaCl, 10 g/L peptone, and 5 g/L yeast extract) supplemented with 50 μg/mL kanamycin. Cells were grown overnight at 250 rpm, 37 °C. Plasmids were purified using the QIAprep Spin Miniprep Kit (Qiagen) according to the manufacturer’s instructions. All cloning results were confirmed by Sanger sequencing (Azenta Life Sciences). An overview of the DNA and protein sequences can be found in the supplementary information.

### Protein expression and purification

#### De novo binder screening constructs

Plasmids encoding the *de novo* binder constructs were transformed into chemically competent *E. coli* BL21 (DE3) and cultured in auto-induction LB medium supplemented with 50 μg/mL kanamycin. Cultures were incubated for 3 hours at 140 rpm, 37 °C, before lowering the temperature to 18 °C for overnight expression. Subsequently, cells were harvested by centrifugation (10 min, 10,000xg, 4 °C) and chemically lysed using Bugbuster protein extraction reagent (Novagen), supplemented with Benzonase endonuclease (Novagen). Lysates were centrifuged at 40,000xg for 30 minutes at 4 °C. Small fractions of cleared supernatant were stored at -70 °C for future assays while the rest was loaded twice onto an equilibrated Ni^2+^-NTA affinity column. Columns were washed with 10 column volumes (CV) of buffer A (50 mM Tris pH 7.4, 370 mM NaCl, 10% (v/v) glycerol, 10 mM imidazole). Proteins were eluted using 3 CV elution buffer (buffer A + 250 mM imidazole). SDS-PAGE was performed to determine protein purity. Correct protein mass was confirmed by Q-ToF LC-MS (WatersMassLynx v4.1), using MagTran v1.03 for MS deconvolution. Mass spectra were obtained using a 1 µL injection volume, containing 0.05 mg/mL of protein in Q-ToF buffer (MilliQ, 0.1% formic acid). Purified proteins were stored in protein storage buffer (50 mM Tris pH 7.4, 150 mM NaCl, 10% glycerol) at -70 °C.

#### LUCOS adapter proteins

The respective plasmids encoding the LUCOS adapters were co-transformed in *E. coli* BL21 (DE3) together with a pEVOL vector encoding a tRNA/ tRNA synthetase pair for the incorporation of the unnatural amino acid pBpA at the amber stop codon^32^. Cells were cultured in LB medium supplemented with 50 µg/mL kanamycin and 25 µg/mL chloramphenicol. At OD_600_ = 0.3, pBpA was added to 1 mM (Bachem, 104504-45-2). Subsequently, cultures were left to grow until OD_600_ = 0.6 at which point protein expression was induced using 1 mM isopropyl β-D-1-thiogalactopyramoside (IPTG) and 0.2% L-arabinose. After overnight expression at 18 °C, cells were harvested by centrifugation (10 min, 10,000xg, 4 °C), followed by resuspension in pre-chilled lysis buffer (50 mM Tris pH 7.4, 500 mM NaCl, 1 mM TCEP, 15 mL/g pellet). Lysis was performed through sonication on ice (15 sec duty cycle at 50% amplitude, Qsonica Q500). Lysates were centrifuged at 40,000xg for 30 minutes at 4 °C. Cleared supernatant was loaded twice onto an equilibrated Ni^2+^-NTA affinity column. Columns were washed with 15 CV of buffer A. Proteins were eluted using 7 CV elution buffer, followed by Strep-Tactin purification according to the manufacturer’s instructions (IBA Life Sciences). The purity of the proteins was determined using SDS-PAGE analysis and the incorporation of pBpA was confirmed by Q-ToF LC-MS (WatersMassLynx v4.1), using MagTran v1.03 for MS deconvolution. Purified proteins were stored in protein storage buffer at -70 °C.

### Protein characterization using circular dichroism (CD) spectroscopy

The secondary structure content was evaluated by CD. Samples were buffer exchanged to Milli-Q and diluted to a final concentration of 0.1 mg/mL. Measurements were performed using the J-815 CD spectrometer (Jasco). Spectra were acquired over a wavelength range of 260 to 190 nm, with a data pitch of 1.0 nm, scan speed of 100 nm/min, response time (DIT) of 0.5 s and bandwidth of 2 nm. Per sample, 5 wavelength scans were collected and averaged.

### Photoconjugation

Monoclonal mouse IgG2a anti-digoxin antibody (26G10) was purchased from ProteoGenix. For the screening assay, pG _LBit^30^ and anti-digoxin antibody were mixed in a 2:1 molar ratio in PBS (pH 7.4) in a final volume of 20 µL and pre-incubated for 30 minutes at room temperature. Photoconjugation was subsequently performed through irradiation with UV light (Thorlabs M365LP1 with a Thorlabs LEDD1B T-Cube LED Driver, λ = 365 nm) for 30 minutes at room temperature. After conjugation, the products were not further purified and stored at 4 °C until use. Photoconjugation with the LUCOS adapter proteins followed the same setup, but with a equimolar mixture of the adapter protein and anti-digoxin antibody. Success of coupling reaction was checked using nonreducing SDS-PAGE analysis.

### Bioluminescent screenings assay

Screening assays were performed with 1 nM of anti-digoxin:pG_LBit and 2.5 µM purified de novo binder or 1 µL of cleared supernatant in a total volume of 20 µL PBS buffer (pH 7.4, 0.1% (w/v) BSA). Triplicate reactions were assembled in NUNC® white 384 flat well plates, briefly spun down and incubated for 30 – 60 minutes at room temperature. Subsequently, 2 µL NLuc substrate furimazine (Promega, N1110) was added to a final 1,000-fold dilution and the bioluminescent emission at 458 nm was recorded in a plate reader (Tecan Spark 10M), with an 25 nm bandwidth and integration time of 100 ms.

### Bioluminescent titrations

*Binder titrations:* Titrations with the *de novo* binders were performed at a constant anti-digoxin:pG_LBit concentration of 1 nM in a total volume of 20 µL PBS buffer in a NUNC® white 384 flat well plate. Binders were diluted to a concentration range of 9.8 nM – 2.5 µM. After incubation of triplicate reactions of the photoconjugated antibody with the binders for 30 – 60 minutes at room temperature, substrate was added (1,000-fold diluted) and bioluminescent emission at 458 nm was recorded in a plate reader (Tecan Spark 10M), with an 25 nm bandwidth and integration time of 100 ms. Response curves were fitted in Origin (2023b).

#### Competitive titrations with digoxin

Digoxin was purchased from Sigma Aldrich (#D6003) and dissolved in DMSO to a final concentration of 10 mM. 500 pM of photoconjugated antibody was incubated with the *de novo* binder in a total volume of 20 µL PBS buffer in NUNC® white 384 flat well plates. Binders were used at a concentration of approximately 0.1 * apparent *K*_*D*_ from the binder titration assays. Next, digoxin was added in a concentration range of 0.5 – 128 nM. Reactions were incubated for an additional 30 minutes – 2 hours, prior to substrate addition (1,000-fold diluted) and bioluminescent emission recordings at 458 nm (25 nm bandwidth, 100 ms integration time). Response curves were fitted in Origin (2023b).

### LUCOS assays

Performance of binders B2 and B6 in the LUCOS sensor was assessed in 20 µL reaction mixtures containing 1 nM LUCOS in PBS buffer, combined with 100 pM split calibrator luciferase, and digoxin diluted to a concentration range of 1.5 nM – 10 µM. Triplicate reactions were assembled in NUNC® white 384 flat well plates, briefly spun down and incubated for 2 hours at room temperature. Subsequently, substrate was added to the reactions to a final 1,000-fold dilution followed by bioluminescent emission recordings at 458 nm (25 nm bandwidth, 100 ms integration time). The blue/green ratio was calculated by dividing bioluminescent emission at 533 nm by emission at 458 nm. Response curves were fitted in Origin (2023b).

## Acknowledgements

We would like to thank Aerin Bogaars, Seth van den Hurk and Tom Philips for performing initial, explorative experiments for this work. We would like to thank Harm van der Veer, Wouter Langers and TU/e Supercomputing Center for their help with installing and running BindCraft. This work was supported by the Dutch research council | Nationaal Regieorgaan Praktijkgericht Onderzoek SIA (RAAK.PRO04.063), NWO small-compute grants EINF-12631 and EINF-13287.

Molecular graphics and analyses performed with UCSF ChimeraX, developed by the Resource for Biocomputing, Visualization, and Informatics at the University of California, San Francisco, with support from National Institutes of Health R01-GM129325 and the Office of Cyber Infrastructure and Computational Biology, National Institute of Allergy and Infectious Diseases.

## Author contributions

Y.d.S. and T.G. conceived and designed the study, performed experiments, analyzed data and wrote the manuscript. M.M. conceived, designed and supervised the study, analyzed the data and wrote the manuscript. All authors discussed the results and commented on the manuscript.

## Data availability

The data that support the plots within this paper and other findings of this study are available from the corresponding author upon reasonable request.

