## supporting information for "*De novo* design of protein competitors for small molecule immunosensing"

**This PDF file includes:**

Figures S1 to S15
Tables S1 and S2
DNA and protein sequences

Figure S12 | Non-reducing SDS-PAGE analysis of photoconjugation of LUCOS_B2 adapter protein 12

Figure S13 | Non-reducing SDS-PAGE analysis of photoconjugation of LUCOS_B6 adapter protein 12

### Supplementary Figures

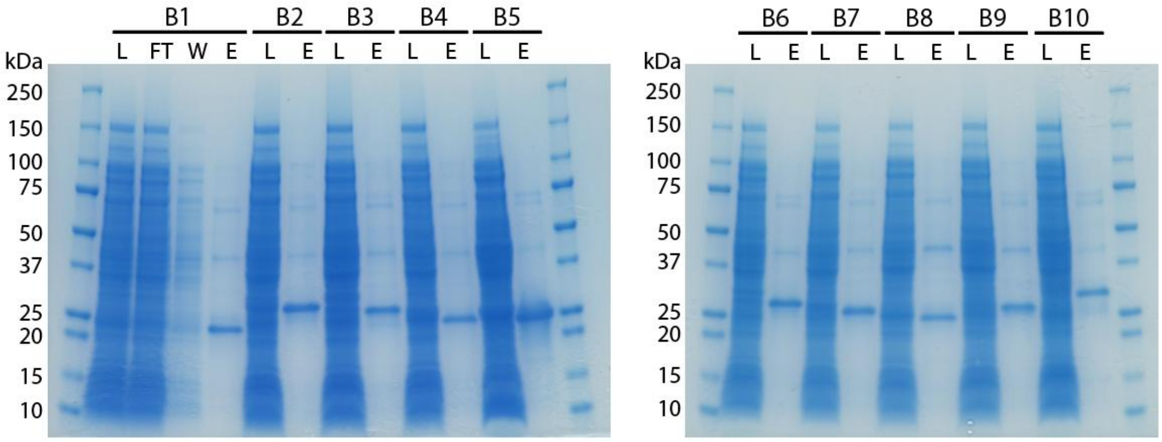

**Figure S1 | SDS-PAGE analysis of expression and purification of *de novo* binders**. *De novo* binders were expressed in *E. coli* and purified using $Ni^{2+}$ affinity chromatography. L: cleared supernatant lysate, FT: flow through, W: wash, E: elution. Marker: Precision Plus Protein™ marker (BioRad). Expected molecular weight binders: B1 = 21 kDa, B2 = 22 kDa, B3 = 23 kDa, B4 = 20 kDa, B5 = 21 kDa, B6 = 23 kDa, B7 = 21.2 kDa, B8 = 20 kDa, B9 = 23 kDa, B10 = 23 kDa.

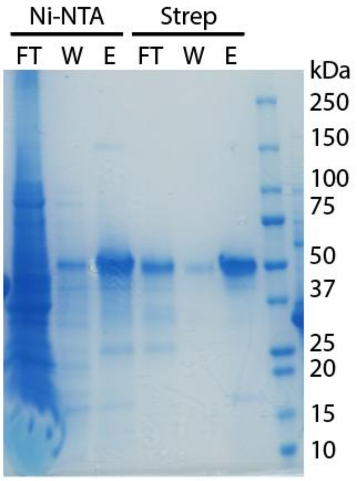

**Figure S2 | SDS-PAGE analysis of expression and purification of pG_dimer_LBit**. Protein G_dimer_LBit construct was expressed in *E. coli* and purified using $Ni^{2+}$ affinity chromatography followed by Strep-Tactin purification. FT: flow through, W: wash, E: elution. Marker: Precision Plus Protein™ marker (BioRad). Expected molecular weight pG_d_LBit = 46 kDa.

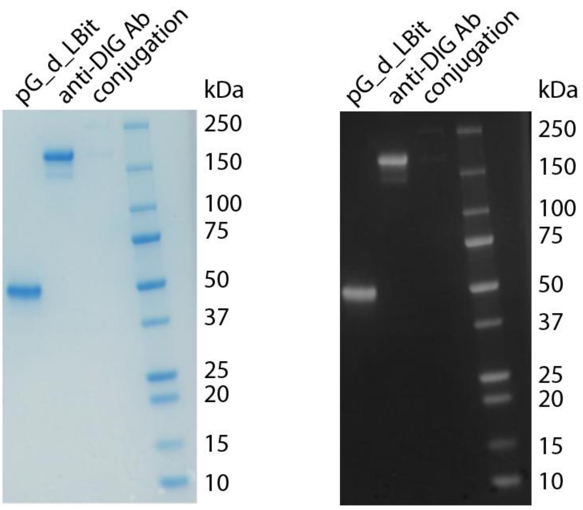

**Figure S3 | Non-reducing SDS-PAGE analysis of photoconjugation of pG_dimer_LBit to anti-DIG antibody**. Purified pG_dimer_LBit was photoconjugated to the anti-digoxin antibody. Mixture containing 1 μM antibody and 2 μM pG_d_LBit was irradiated for 30 minutes with UV light (λ = 365 nm). Marker: Precision Plus Protein™ marker (BioRad). Expected molecular weight pG_d_LBit = 46 kDa, anti-DIG Ab = 150 kDa, conjugation product = 196 kDa. Left: original image, right: color inverted and increased contrast of original image using Adobe Photoshop 2026 software.

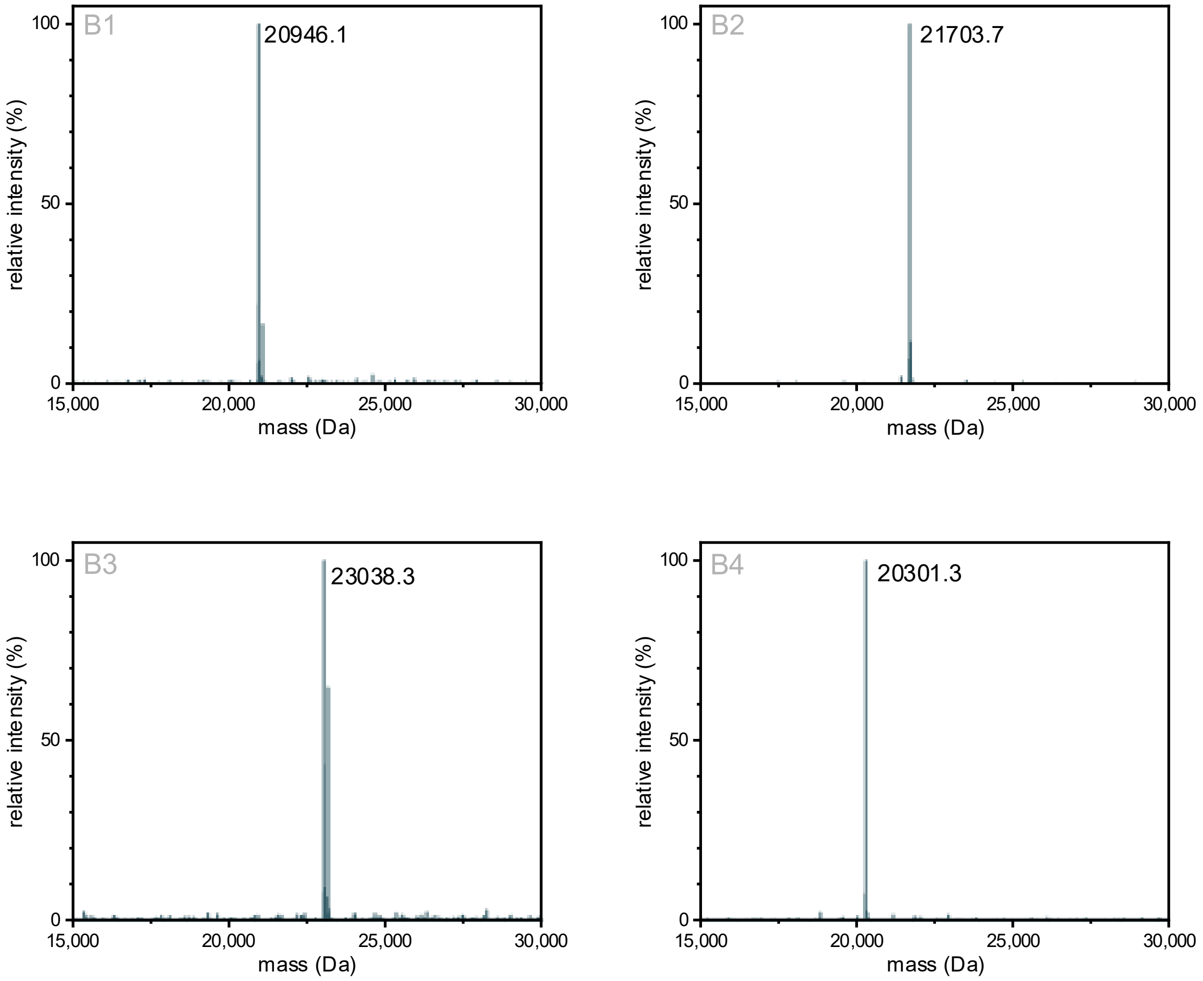

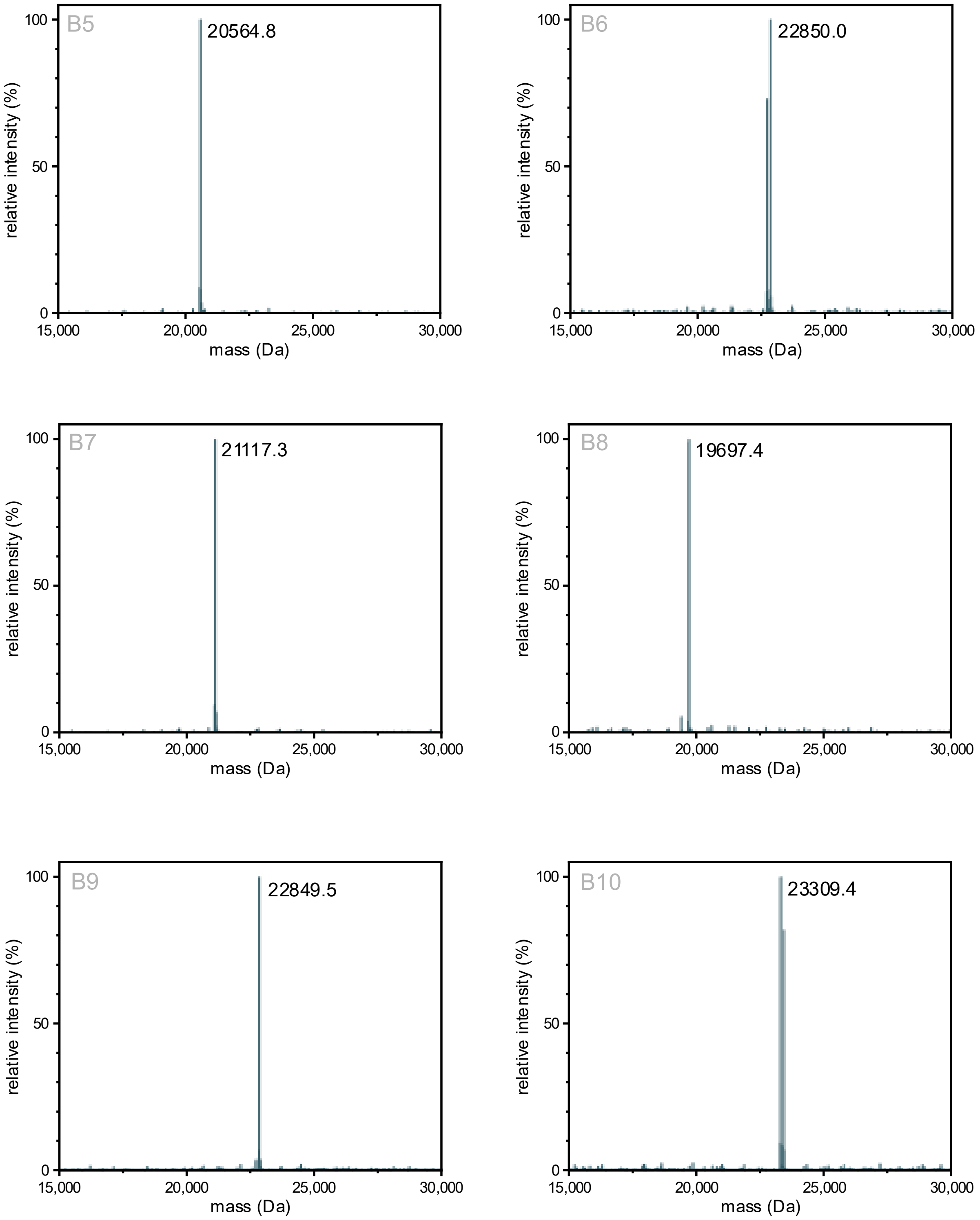

**Figure S4 | ESI-QTOF mass spectra of *de novo* binders B1-B10**. Theoretical masses: B1 = 21077.4 Da (N-terminal methionine excision (NME): 20946.2 Da), B2 = 21834.9 Da (NME: 21703.7 Da), B3 = 23169.7 Da (NME: 23038.5 Da), B4 = 20432.5 Da (NME: 20301.3 Da), B5 = 20696.0 Da (NME: 20564.8 Da), B6 = 22850.1 Da, B7 = 21248.5 Da (NME: 21117.3 Da), B8 = 19828.8 Da (NME: 19697.6 Da), B9 = 22849.5 Da, B10 = 23440.7 Da (NME: 23309.5 Da).

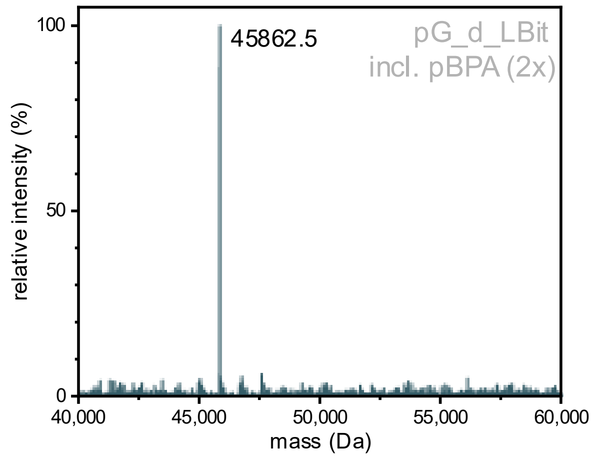

**Figure S5 | ESI-QTOF mass spectra of pG_dimer_LB**. Theoretical mass pG_dimer_LB including 2X pBpA = 45994.2 Da (N-terminal methionine excision: 45863.0 Da).

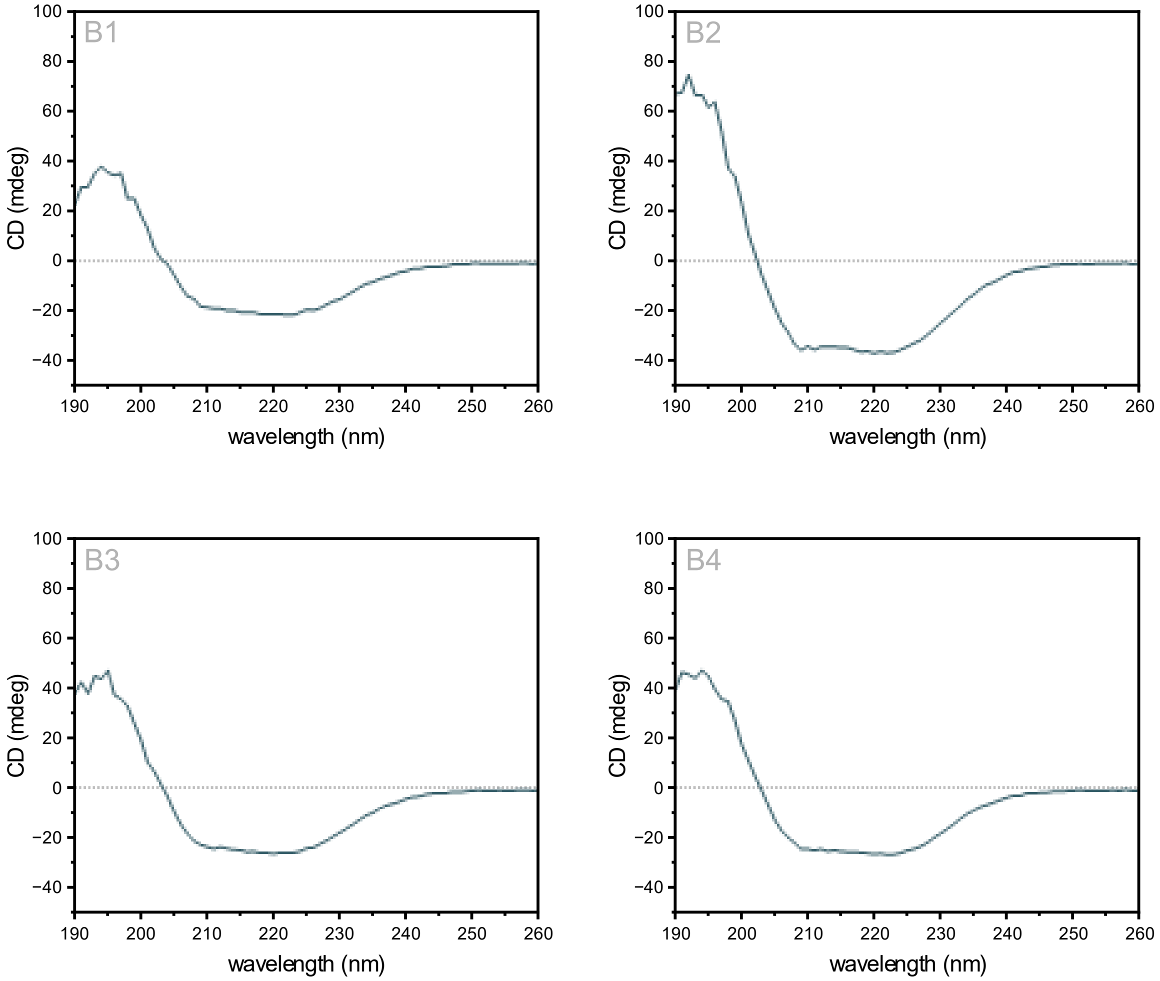

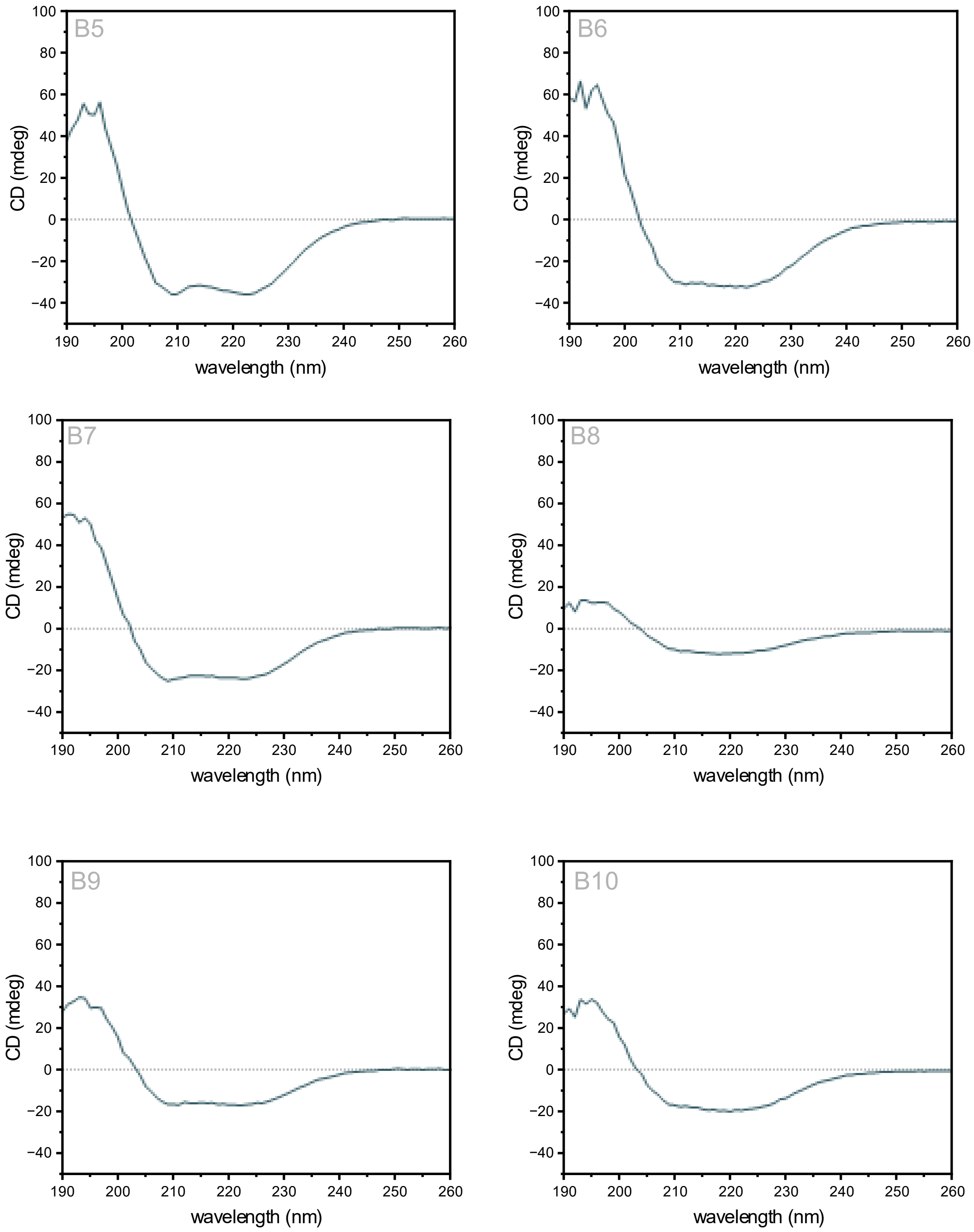

**Figure S6 | Circular dichroism *de novo* binders.** CD confirms the presence of the predicted α-helical secondary structure of the *de novo* designed binders.

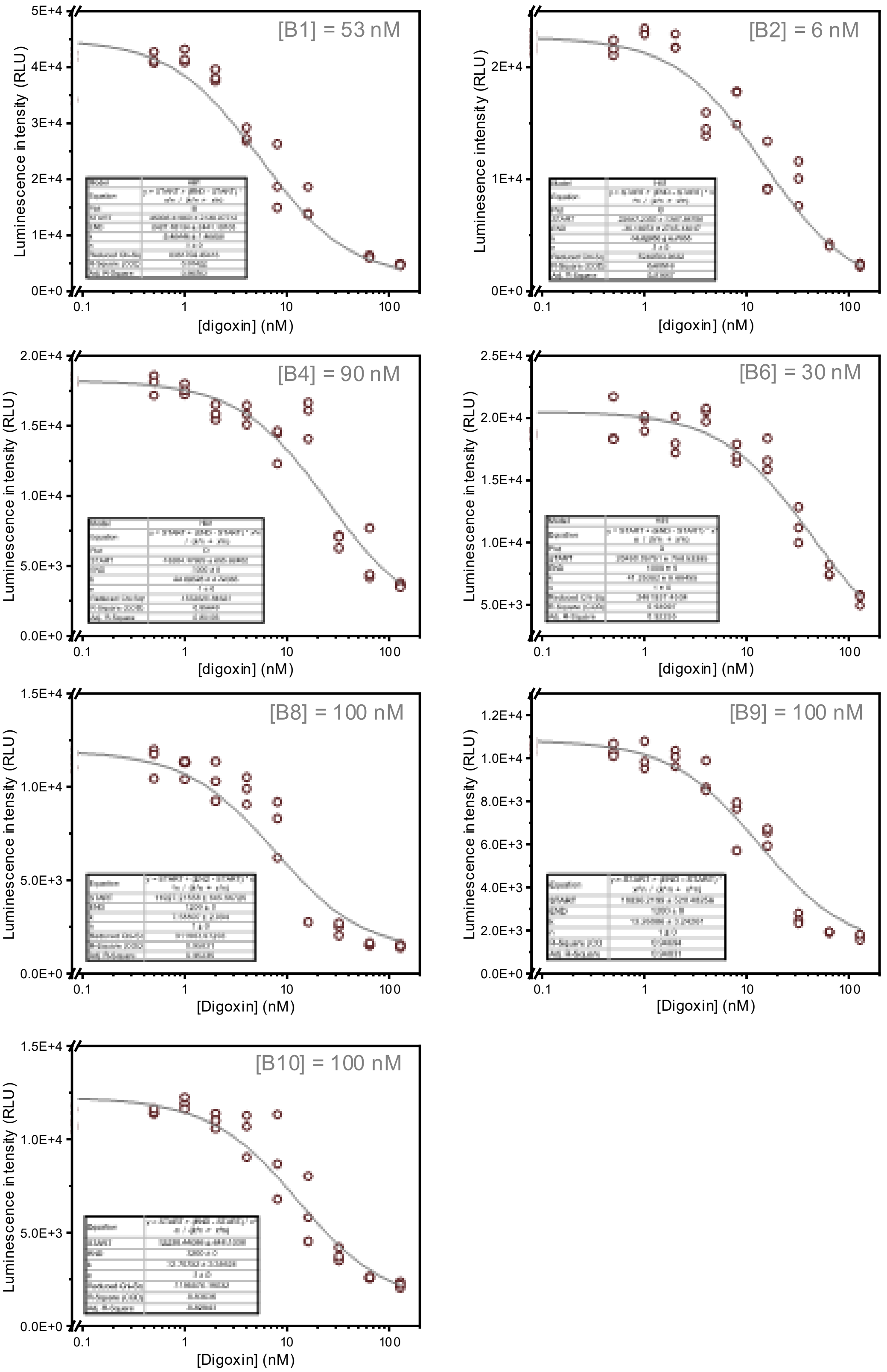

**Figure S7 | Luminescent competition assay *de novo* binders**. Competitive binding was confirmed by titrated digoxin ($K_D$ = 9 nM) to the formed binder-Ab complexes. Reactions were incubated for 2 hours prior to luminescence measurement to ensure that equilibrium was reached. Anti-DIG:LB conjugation was used at a constant concentration of 1 nM, and binder concentrations were used as indicated in the figures. The acquired data was fitted with a Hill function with offset using Origin2023b software. Insets represent the fitting parameters and adjusted R-squared value. For all binders, the Michaelis Menten constant (n) was fixed to 1 assuming no cooperativity. Circles represent individual data points and lines represent model fits.

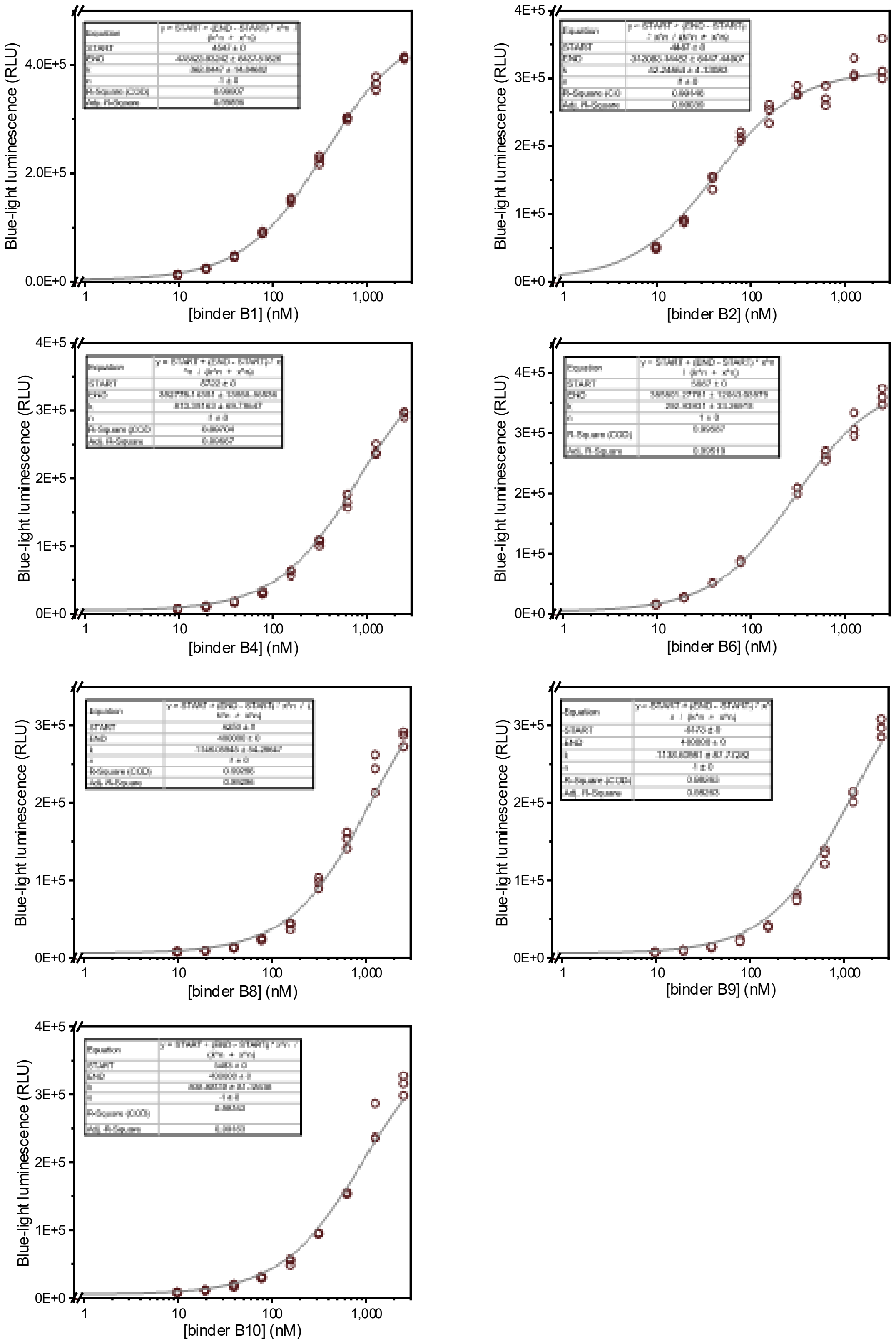

**Figure S8 | Luminescent binding assays *de novo* binders**. Binding of *de novo* proteins to the anti-DIG antibody was confirmed by titrating the binder_SBit constructs (8.9 nM – 2.5 µM) against a constant concentration of anti-DIG_LBit (1 nM). Reactions were incubated for 30 minutes at room temperature prior to luminescence measurement to ensure that equilibrium was reached. The acquired data was fitted with a Hill function with offset using Origin2023b software. Insets represent the fitting parameters and adjusted R-squared value. For all binders, the Michaelis Menten constant (n) was fixed to 1 assuming no cooperativity. Circles represent individual data points and lines represent model fits.

### Thermodynamic model

To model the LUCOS response in presence of *de novo* competitors with different affinities, we adapted the thermodynamic model described in *van Aalen et al*. (2024). This model defines the equilibrium equations involved in the different binding states of the LUCOS sensor (Figure S9A) to simulate the sensor response to increasing concentrations of target analyte. Baseline parameters were kept consistent with the original model, except for the antibody-analyte affinity (9 nM, for digoxin) and the affinity of the competitor domain (42 nM, in Figure S9B and 293 nM in Figure S9C). Although effective molarities of competitor and SBiT domains are likely altered due to the addition of a protein-based competitor rather than a small-molecule analog, these values are hard to model precisely and are not strictly necessary for a relative comparison between the different affinity competitors.

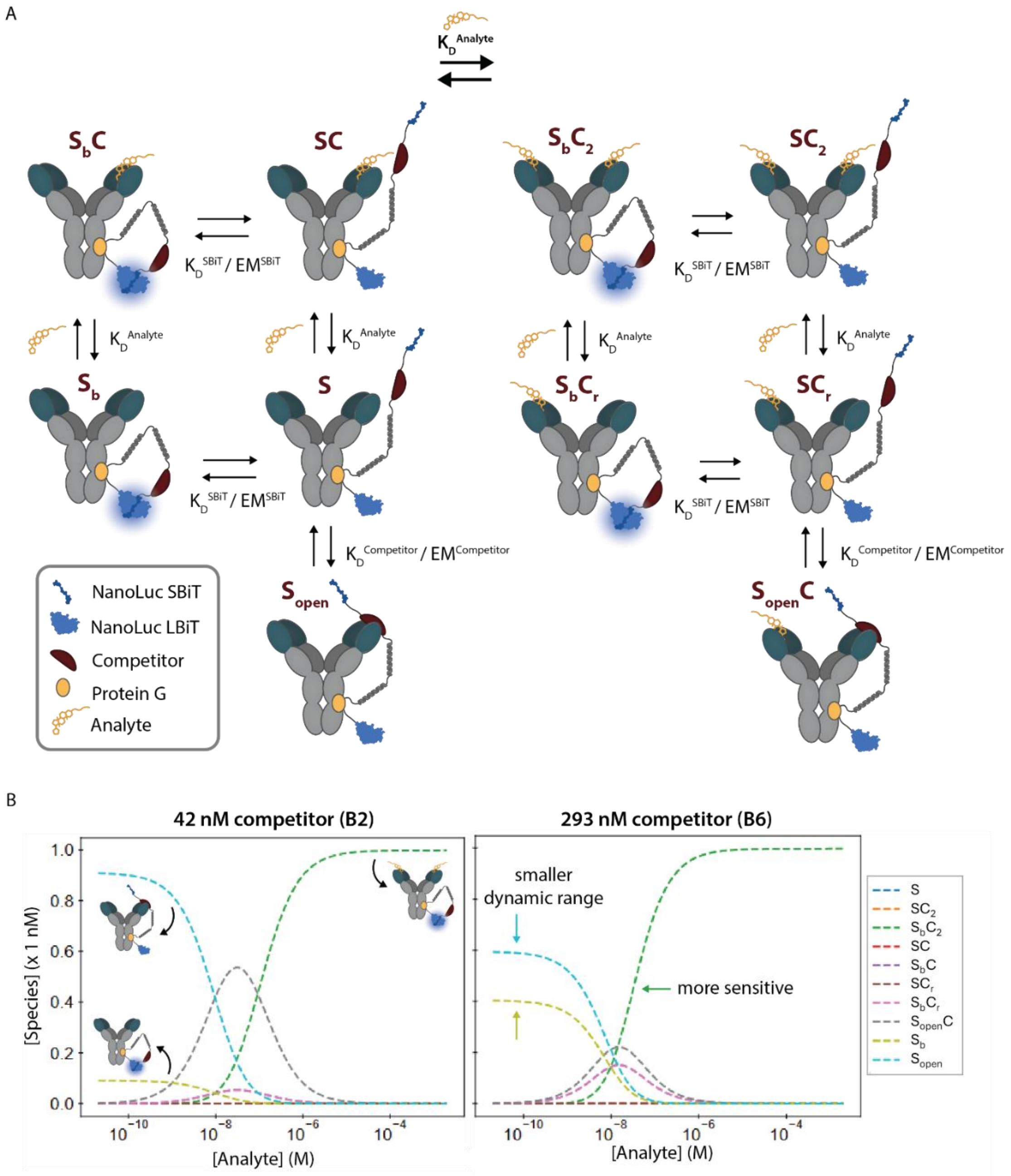

Figure S9 | LUCOS thermodynamic model A) Schematic of different sensor states. B) Modelled LUCOS response to increasing concentrations of analyte (digoxin) using competitors B2 and B6. Baseline parameters: $[S_{tot}]$ = 1 nM, $K_{D,Analyte}$ = 9 nM, $K_{D,SBiT}$ = 2.5 μM, $EM_{competitor}$ = 191 μM and $EM_{SBiT}$ = 1.08 mM.

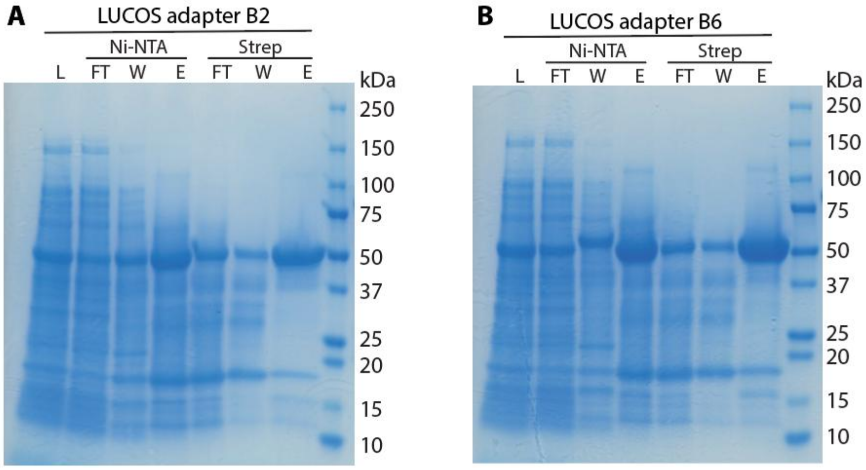

**Figure S10 | SDS-PAGE analysis of expression and purification LUCOS adapter proteins**. LUCOS adapter proteins were expressed in *E. coli* and purified using $Ni^{2+}$ affinity chromatography followed by Strep-Tactin purification. L: cleared supernatant lysate, FT: flow through, W: wash, E: elution. Marker: Precision Plus Protein™ marker (BioRad). (A) Expected molecular weight LUCOS_B2 = 54 kDa. (B) Expected molecular weight LUCOS_B6 = 55 kDa.

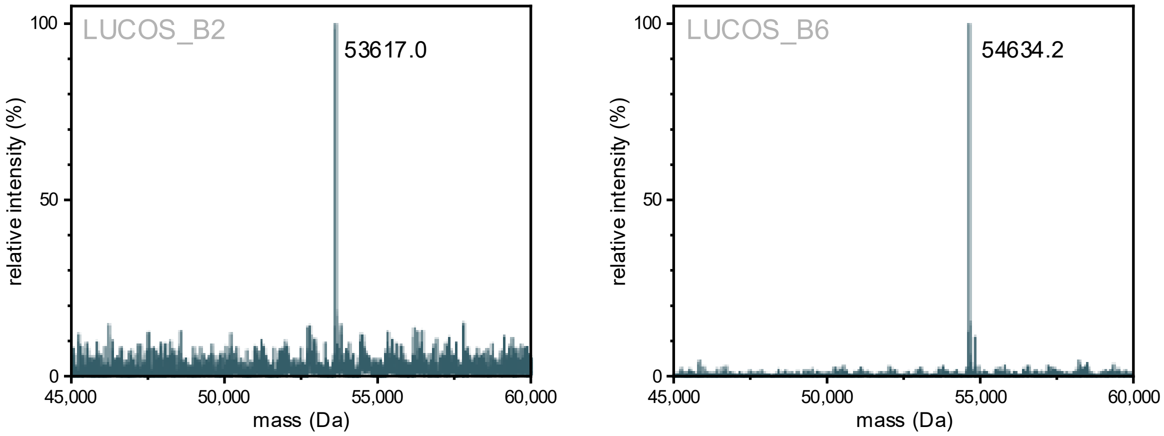

**Figure S11 | ESI-QTOF mass spectra of LUCOS adapter proteins**. Theoretical masses: LUCOS_B2 (incl. 1X pBpA) = 53750.1 Da (N-terminal methionine excision (NME): 53618.9 Da), LUCOS_B6 (incl. 1X pBpA) = 54765.2 Da (NME: 54634.0 Da).

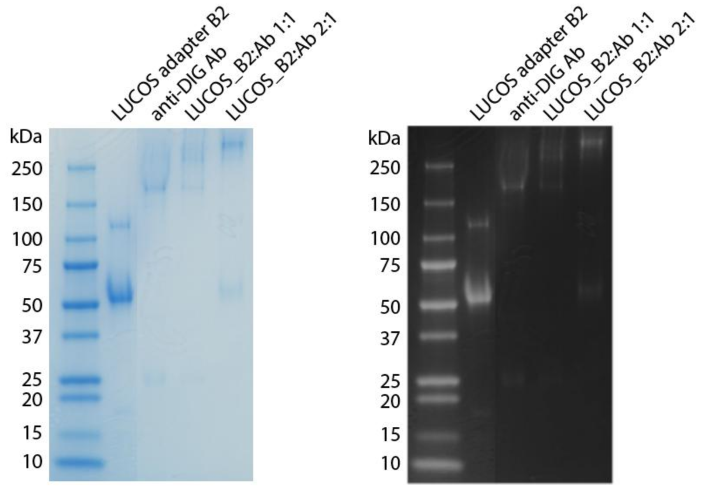

**Figure S12 | Non-reducing SDS-PAGE analysis of photoconjugation of LUCOS_B2 adapter protein**. Purified LUCOS_B2 adapter was photoconjugated to the anti-digoxin antibody. Mixture containing 1 µM antibody and 1 µM of 2 µM LUCOS_B2 adapter was irradiated for 30 minutes with UV light (λ = 365 nm). Marker: Precision Plus Protein™ marker (BioRad). Expected molecular weight LUCOS_B2 = 54 kDa, anti-DIG Ab = 150 kDa, conjugation product = 194 kDa. Left: original image, right: color inverted and increased contrast of original image using Adobe Photoshop 2026 software.

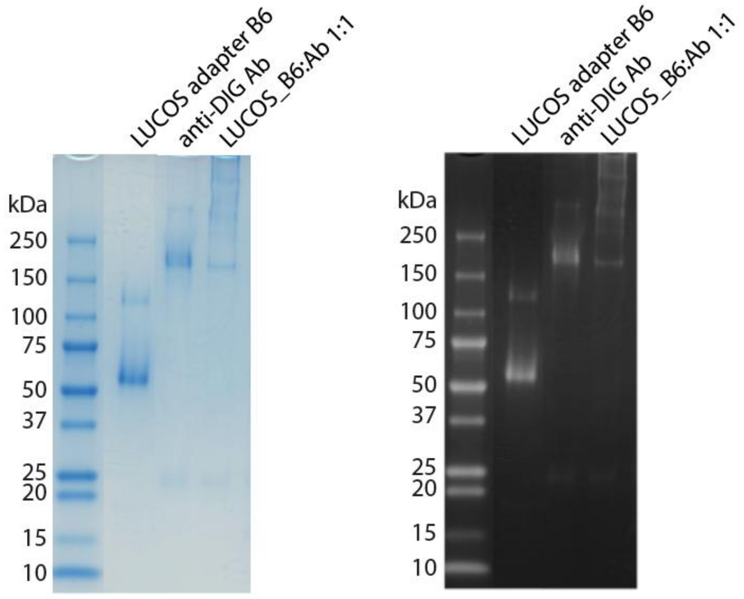

**Figure S13 | Non-reducing SDS-PAGE analysis of photoconjugation of LUCOS_B6 adapter protein**. Purified LUCOS_B6 adapter was photoconjugated to the anti-digoxin antibody. Mixture containing 2 µM antibody and LUCOS_B6 adapter was irradiated for 30 minutes with UV light (λ = 365 nm). Marker: Precision Plus Protein™ marker (BioRad). Expected molecular weight LUCOS_B2 = 55 kDa, anti-DIG Ab = 150 kDa, conjugation product = 195 kDa. Left: original image, right: color inverted and increased contrast of original image using Adobe Photoshop 2026 software.

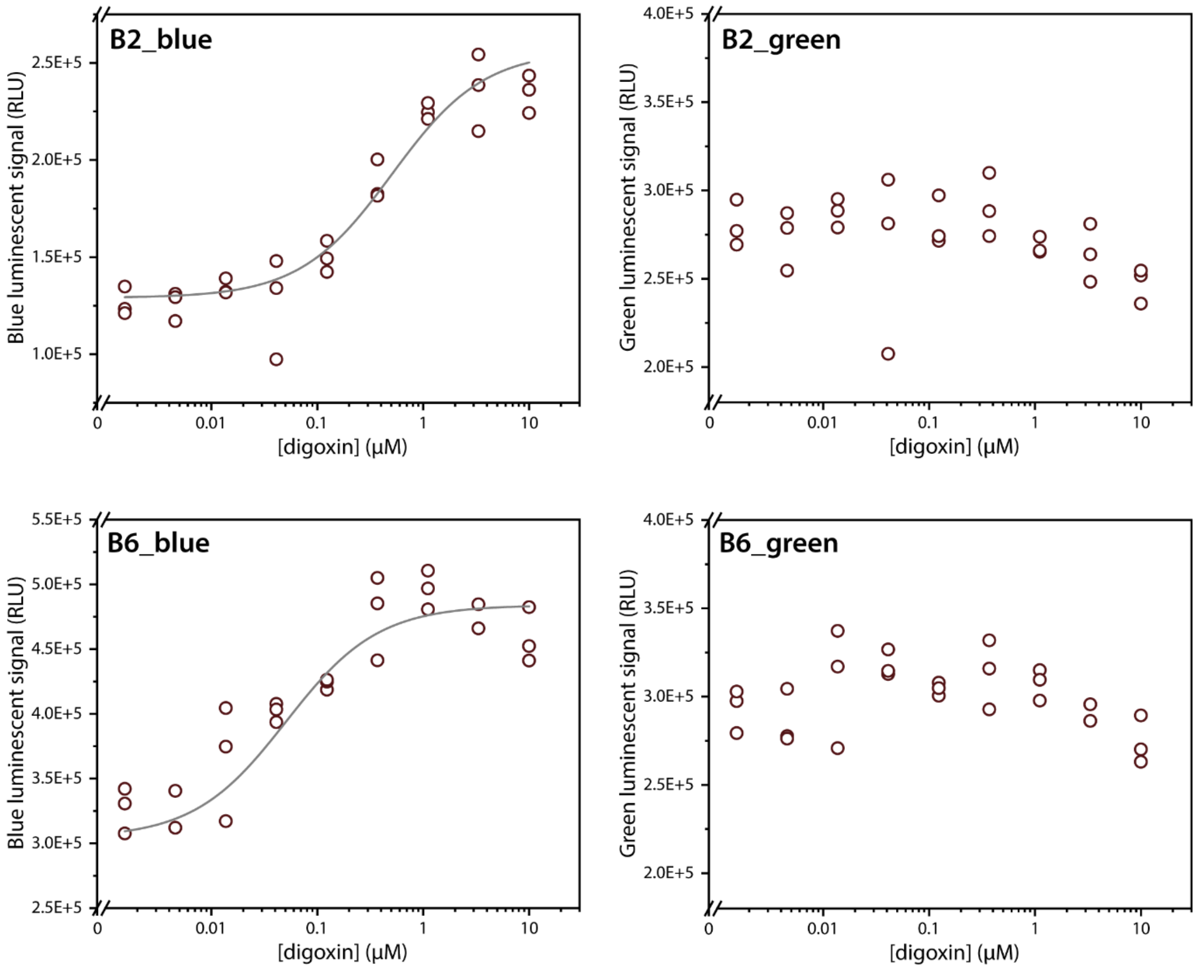

**Figure S14 | Absolute blue and green signals LUCOS_B2 and _B6**. Luminescent signals in the blue (λ = 458 nm) and green (λ = 533 nm) channel of the LUCOS_B2 and LUCOS_B6 sensors. 1 nM LUCOS sensor and 100 pM calibrator was incubated with 1.5 nM – 10 μM digoxin for 2 hours prior to luminescent readout. Circles represent individual data points and lines represent model fits.

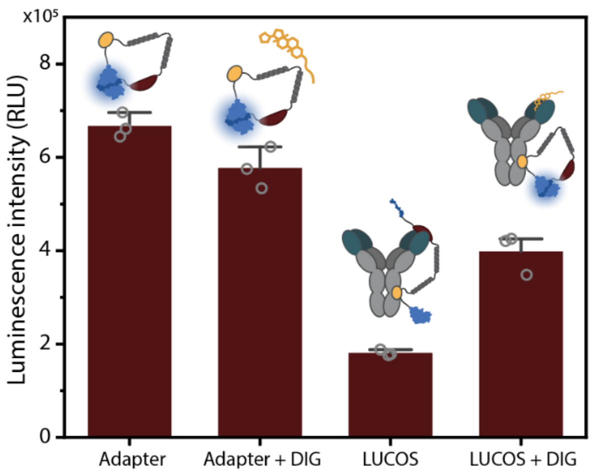

**Figure S15 | LUCOS_B2 bioluminescent background signal and response**. Luminescent signal of LUCOS_B2 adaptor protein, LUCOS conjugation, and effect of digoxin were screened in a bioluminescent assay. Adapter = 1 nM LUCOS_B2 adapter protein, Adapter + DIG = 1 nM LUCOS_B2 adapter protein + 10 μM digoxin, LUCOS = 1 nM photoconjugated LUCOS_B2, LUCOS + DIG = 1 nM photoconjugated LUCOS_B2 + 10 μM digoxin. Samples were assembled in 20 μL 1X PBS + 0.1% (w/v) BSA, and incubated at room temperature for 30 minutes prior to readout of blue luminescent signal (λ = 458 nm). Individual data points (n = 3) are represented as circles.

### Supplementary Tables

**Table S1 | Hotspots and confidence metrics *de novo* binders.**

| Binder | Hotspot | AF3 iPTM | AlphaBridge contact iPTM |
|---|---|---|---|
| B1 | W103 | 0.91 | 0.79-0.87 |
| B2 | W103 | 0.93 | 0.86-0.88 |
| B3 | W103 | 0.88 | 0.82-0.85 |
| B4 | Y32 | 0.91 | 0.82-0.86 |
| B5 | Y32 | 0.87 | 0.71-0.76 |
| B6 | Y32 | 0.89 | 0.79-0.83 |
| B7 | Y32 | 0.89 | 0.73-0.78 |
| B8 | V99 | 0.92 | 0.85-0.87 |
| B9 | V99 | 0.90 | 0.83 |
| B10 | V99 | 0.91 | 0.86 |

**Table S2 | Cloning primers.**

| Primer | Description | Sequence |
|---|---|---|
| YS_130 | FOR backbone linearization *de novo*-SBit | GAATTTACAGGAGGTTCGGGTG |
| YS_131 | REV backbone linearization *de novo*-SBit | GCCCATGGTATATCTCCTTCTTAAAG |
| YS_132 | FOR AgeI overhang addition to binders | AGTCACCGGTCGACCCACCCGAACCTCCTGTAAATTC |
| YS_133 | REV SpeI overhang addition to binder B2 | AGTGACTAGTATGGGCAAAAAATCAGACGAAGAGGTCG |
| YS_134 | REV SpeI overhang addition to binder B6 | AGTGACTAGTATGGGCAGTAAGGAAGAAGAAGTCGAACG |

### Protein coding sequences and translations

#### Protein sequence anti-digoxin variable domains used for competitor design

Truncated anti-digoxin ScFv protein sequence used for AlphaFold2 prediction as target input for the BindCraft pipeline. Color codes: anti-DIG heavy chain highlighted in yellow; anti-DIG light chain highlighted in green.

VQLQQSGPELVKPGASVRMSCKSSGYIFTDFYMNWVRQSHGKSLDYIGYISPYSGVTGYNQKFKGKATLTVDKSSSTAY
MELRSLTSEDSAVYYCAGSSGNKWAMDYWGHGASVTV

DVVMTQTPLSLPVSLGDQASISCRSSQSLVHSNGNTYLNWYLQKAGQSPKLLIYKVSNRFSGVPDRFSGSGSGTDFTLK
ISRVEAEDLGIYFCSQTTHVPPTFGGGTKLEIK

#### pG_dimer_LBit

Color codes: Strep-Tag II sequence highlighted in red; protein G dimer highlighted in yellow; amber codon (overridden for pBpA insertion) highlighted in green; semi-flexible linker highlighted in pink; LBit highlighted in blue; 6xHis-tag highlighted in gray.

```
ATGGGCTGGAGCCATCCGCAGTTTGAAAAAGGTGGTAGCATGACATTTAAACTGATTATCAACGGCAAAACTTTAAAG
 M  G  W  S  H  P  Q  F  E  K  G  G  S  M  T  F  K  L  I  I  N  G  K  T  L  K

GGAGAGATCACAATAGAAGCGGTGGATGCTTAGGAGGCGGAGAAGATTTTTAAGCAGTATGCAAATGATTATGGAATT
 G  E  I  T  I  E  A  V  D  A  *  E  A  E  K  I  F  K  Q  Y  A  N  D  Y  G  I

GATGGTGAATGGACTTATGACGACGCAACTAAAACTTTCACGGTAACAGAAGAATTTACAGGAGGTTCGGGTGGGTCG
 D  G  E  W  T  Y  D  D  A  T  K  T  F  T  V  T  E  E  F  T  G  G  S  G  G  S

GGAGGTTCTGGCGGCTCTGGAGGAAGTGGTGGTAGCGGTGAATTCGCCGAAGCAGCCGCTAAAGAAGCCGCAGCAAAG
 G  G  S  G  G  S  G  G  S  G  G  S  G  E  F  A  E  A  A  A  K  E  A  A  A  K

GAAGCCGCGGCCAAGGAGGCAGCCGCAAAAGAGGCCGCGGCGAAGGAAGCAGCAGCCAAGGCAGAATTCGGGGGTAGC
 E  A  A  A  K  E  A  A  A  K  E  A  A  A  K  E  A  A  A  K  A  E  F  G  G  S

GGCGGCTCGGGGGGTAGTGGTGGAAGCGGGGGTTCAGGCGGTTCTGGGGGTACCATGACATTTAAACTGATAATCAAC
 G  G  S  G  G  S  G  G  S  G  G  S  G  G  S  G  G  T  M  T  F  K  L  I  I  N

GGCAAAACCTTAAAAGGGGAGATCACAATTGAGGCAGTCGATGCCTAGGAAGCCGAGAAAATCTTTAAACAATATGCT
 G  K  T  L  K  G  E  I  T  I  E  A  V  D  A  *  E  A  E  K  I  F  K  Q  Y  A

AATGATTATGGTATTGACGGAGAATGGACGTATGACGATGCGACAAAAACTTTCACCGTAACTGAGCTCACAGGAGGT
 N  D  Y  G  I  D  G  E  W  T  Y  D  D  A  T  K  T  F  T  V  T  E  L  T  G  G

TCGGGTGGGTCGGGAGGTTCTGGCGGCTCTGGAGGAAGTGGTGGTAGCGGTGAATTCGCCGAAGCAGCCGCTAAAGAA
 S  G  G  S  G  G  S  G  G  S  G  G  S  G  G  S  G  E  F  A  E  A  A  A  K  E

GCCGCAGCAAAGGAAGCCGCGGCCAAGGAGGCAGCCGCAAAAGAGGCCGCGGCGAAGGAAGCAGCAGCCAAGGCAGAA
 A  A  A  K  E  A  A  A  K  E  A  A  A  K  E  A  A  A  K  E  A  A  A  K  A  E

TTCGGGGGTAGCGGCGGCTCGGGGGGTAGTGGTGGAAGCGGGGGTTCAGGCGGTTCTGGGGGTACCGTCTTCACACTC
 F  G  G  S  G  G  S  G  G  S  G  G  S  G  G  S  G  G  T  V  F  T  L
```

GAAGATTTCGTTGGGGACTGGGAACAGACAGCCGCCTACAACCTGGACCAAGTCCTTGAACAGGGAGGTGTGTCCAGT
E D F V G D W E Q T A A Y N L D Q V L E Q G G V S S

TTGCTGCAGAATCTCGCCGTGTCCGTAACTCCGATCCAAAGGATTGTCCGGAGCGGTGAAAATGCCCTGAAGATCGAC
L L Q N L A V S V T P I Q R I V R S G E N A L K I D

ATCCATGTCATCATCCCGTATGAAGGTCTGAGCGCCGACCAAATGGCCCAGATCGAAGAGGTGTTTAAGGTGGTGTAC
I H V I I P Y E G L S A D Q M A Q I E E V F K V V Y

CCTGTGGATGATCATCACTTTAAGGTGATCCTGCCCTATGGCACACTGGTAATCGACGGGGTTACGCCGAACATGCTG
P V D D H H F K V I L P Y G T L V I D G V T P N M L

AACTATTTCGGACGGCCGTATGAAGGCATCGCCGTGTTCGACGGCAAAAAGATCACTGTAACAGGGACCCTGTGGAAC
N Y F G R P Y E G I A V F D G K K I T V T G T L W N

GGCAACAAAATTATCGACGAGCGCCTGATCACCCCCGACGGCTCCATGCTGTTCCGAGTAACCATCAACAGCGGAGGT
G N K I I D E R L I T P D G S M L F R V T I N S G G

TCCCACCACCACCATCACCAC
S H H H H H H

#### *De novo* binder B1_SBit

Color codes: *de novo* binder highlighted in yellow; semi-flexible linker highlighted in pink; SBit ($K_D$ 190 μM) highlighted in cyan; 6xHis-tag highlighted in gray.

ATGGGCATGCTTAGCCCGGCAGAACTGCTGCAGAAAGTTTTTAGTAAACGCGAACTGCGTCGAATGTTAAGGGATGTT
M G M L S P A E L L Q K V F S K R E L R R M L R D V

TACATGGTTCTGCGATGGTCCGCTACCGAAGAAGAAGCCGTTGAGCAGGTGGAAAAAATGCTGGAGTATGTGCTTAAC
Y M V L R W S A T E E E A V E Q V E K M L E Y V L N

GACCTCAAGGAAAAATATGGTGAGGATAGTGAGCGGGTCAAACTGTTCGAAGAATTTATCAAAAAATTACGGGAGCAA
D L K E K Y G E D S E R V K L F E E F I K K L R E Q

GGAAAAAACGGGCGTATTGACCCTCGTCTGCTGAAAGAAGTGATTGAAGAAGCGATCAGGAATTACCCTGAATTTACA
G K N G R I D P R L L K E V I E E A I R N Y P E F T

GGAGGTTCGGGTGGGTCGGGAGGTTCTGGCGGCTCTGGAGGAAGTGGTGGTAGCGGTGAATTCGCCGAAGCAGCCGCT
G G S G G S G G S G G S G G S G G S G E F A E A A A

AAAGAAGCCGCAGCAAAGGAAGCCGCGGCCAAGGAGGCAGCCGCAAAAGAGGCCGCGGCGAAGGAAGCAGCAGCCAAG
K E A A A K E A A A K E A A A K E A A A K E A A A K

GCAGAATTCGGGGGTAGCGGCGGCTCGGGGGGTAGTGGTGGAAGCGGGGGTTCAGGCGGTTCTgggGGTACCGTTACC
A E F G G S G G S G G S G G S G G S G G S G G T V T

GGCTATCGTCTGTTTGAAGAAATTCTCGGCGGTTCACATCATCATCACCACCAT
G Y R L F E E I L G G S H H H H H H

#### *De novo* binders B2-B10

*De novo* binders B2-B10 were expressed in the same construct as B1 (coupled to SBit $K_D$ = 190 µM). Here, only the binder sequences are given. Color code: *de novo* binder highlighted in yellow.

**Binder B2**:

```
ATGGGCAAAAAATCAGACGAAGAGGTCGGTCTCGAATTAGCTAAAGAGATTCTCAAGGAACTGATTGAATCAATGGGG
 M  G  K  K  S  D  E  E  V  G  L  E  L  A  K  E  I  L  K  E  L  I  E  S  M  G

TTGAGCGAAGTTCCCAAAAAAGAGGAACTGGACCATTTCCTGCCGCACCTGGAATGGGACGCGGACTGGCAGATACAG
 L  S  E  V  P  K  K  E  E  L  D  H  F  L  P  H  L  E  W  D  A  D  W  Q  I  Q

GAACGTATTGTCGAATACTATAAAGAAAACGGAGAAGAACCGACGGAGGAACGCCTTGAAACAGCCCATAAAGCGGCG
 E  R  I  V  E  Y  Y  K  E  N  G  E  E  P  T  E  E  R  L  E  T  A  H  K  A  A

TGGTCTGTACTGCAGAAATTTATCGCGGAAGTCCGTGCCATGGCAGCAAAAGGCGATCCGTCGAAAGAAGAAATTTTG
 W  S  V  L  Q  K  F  I  A  E  V  R  A  M  A  A  K  G  D  P  S  K  E  E  I  L

GAGGTCATTGAATCCCTTGAAGCGGAATTT
 E  V  I  E  S  L  E  A  E  F
```

**Binder B3**:

```
ATGGGCTCAGTGTATGAGGAGATTCGCGAACTGTTAGGGAAAACGCTGCACGATTTTGCTCAAGGTAAACTGAGCAAA
 M  G  S  V  Y  E  E  I  R  E  L  L  G  K  T  L  H  D  F  A  Q  G  K  L  S  K

GAAGAAGTGCTCAAAAAAATCGAGGAAGCAGTGGAAAAGCTTGAAAAATACTTTGAAGAAAATAAAGACGTGCCAGGT
 E  E  V  L  K  K  I  E  E  A  V  E  K  L  E  K  Y  F  E  E  N  K  D  V  P  G

GTGGAGGAATTATACGAAAAATACAAAAAGGAAGTGGAAAAATATATGTCGTGGCTTCGGACCATGCCTTTTGACCCG
 V  E  E  L  Y  E  K  Y  K  K  E  V  E  K  Y  M  S  W  L  R  T  M  P  F  D  P

AGCAATCCTCAGCAGGTCTGGGATTGGCATATGGACTTGATGAGCTTGGGATACTATCTGCGTAAATACGCAAAAGAA
 S  N  P  Q  Q  V  W  D  W  H  M  D  L  M  S  L  G  Y  Y  L  R  K  Y  A  K  E

CTCGAAGAATTGGTGAAAAAAGCTGAAGAACTGGAAAAGCTTGAATTT
 L  E  E  L  V  K  K  A  E  E  L  E  K  L  E  F
```

**Binder B4**:

```
ATGGGCAGTTCAATGGAAGAAGAATTGCGGAAAAAAGTCGAAGAAGCTCGGGAAAAAATGATAGAGCTGATAGGTGAA
 M  G  S  S  M  E  E  E  L  R  K  K  V  E  E  A  R  E  K  M  I  E  L  I  G  E

GAACGGGTAGAATGGATAGAAGATTGGGCGCTGTTTGTCATCCACATGCATAAATGGGCTGGACACACCGATGTCGAG
 E  R  V  E  W  I  E  D  W  A  L  F  V  I  H  M  H  K  W  A  G  H  T  D  V  E

GAACTGGCCGAATTTGTCGTGAAACATGTTGGGGATATTTTTAAGCGCGAGTTCCCCGAGATCCCGGAGGAGGTGCAG
 E  L  A  E  F  V  V  K  H  V  G  D  I  F  K  R  E  F  P  E  I  P  E  E  V  Q

AAAAAGATCCTGGAATTACTCTACGAATATGCCTATGCCCAGGCCAAACTGGAAGAATTT
 K  K  I  L  E  L  L  Y  E  Y  A  Y  A  Q  A  K  L  E  E  F
```

**Binder B5:**

```
ATGGGCATGAAATTAAGTGAGGAAGAGTTCAAGAAAAAAGTGGAAGAACTTGCTAGAGCCTTCATGATGATCGCGTGG
 M  G  M  K  L  S  E  E  E  F  K  K  K  V  E  E  L  A  R  A  F  M  M  I  A  W

GATAAAGCCTGGTTCCCGGAAGTCACCGAAGAGAAAGCTCGTGAAATGTTAAAAGAAGAGTTACGGAAGTACTTTCCA
 D  K  A  W  F  P  E  V  T  E  E  K  A  R  E  M  L  K  E  E  L  R  K  Y  F  P

GAAGCGCCGGAAGAAAAAATTGAGAAATACGTGGAGAAATTCCTTAAAGAGGCAGAAGAGCTGTGGAAACAAGGGAAA
 E  A  P  E  E  K  I  E  K  Y  V  E  K  F  L  K  E  A  E  E  L  W  K  Q  G  K

TTAACGTGGGAGAAAGTGAAAGCGTTTGCAGAAGAAATCGTCAAAGAATTTAATGGCGAATTT
 L  T  W  E  K  V  K  A  F  A  E  E  I  V  K  E  F  N  G  E  F
```

**Binder B6:**

```
ATGGGCAGTAAGGAAGAAGAAGTCGAACGCCACATCTATTTACATCTAAGAGATCTGGACATGATGCTGCATTGGGAA
 M  G  S  K  E  E  E  V  E  R  H  I  Y  L  H  L  R  D  L  D  M  M  L  H  W  E

CAGTTTGACGAGGCCCTGGCGTTCGCGAAAAAATCTCTGGAGGAGCTGGTCAAACTGCTGAAAGGTTTCTCTAAAGAA
 Q  F  D  E  A  L  A  F  A  K  K  S  L  E  E  L  V  K  L  L  K  G  F  S  K  E

GAGTATGAAAAAATCCTGACGGAATACTGGGAACGTGTTGTTGAGTTGATCCGCCGTTACGCCGATTGGTACGGCGCG
 E  Y  E  K  I  L  T  E  Y  W  E  R  V  V  E  L  I  R  R  Y  A  D  W  Y  G  A

GGTGGCAACGGTCGTCGTGACCCGGCCGCCTACCAAGCGACGGCGGACCGTATGATAGCCGAACTGTCTAAAGTATAT
 G  G  N  G  R  R  D  P  A  A  Y  Q  A  T  A  D  R  M  I  A  E  L  S  K  V  Y

AACGACATTCTGGAAAACTACGACAAATTACAAGCGGAGGAATTT
 N  D  I  L  E  N  Y  D  K  L  Q  A  E  E  F
```

**Binder B7:**

```
ATGGGCAGTAAACTGGAAGAAGTCCTTAAGGAGACCAAAAAGATTTTTAGTGAGGCGTACGAAAAAGTGGCGAAAGCT
 M  G  S  K  L  E  E  V  L  K  E  T  K  K  I  F  S  E  A  Y  E  K  V  A  K  A

GCGCCGAAGGACCCCAAAGCCCAACCGCGTTTCGATCAAATGGAAGCGCTGGATAAAACTCTTCGTGAATTTGATGAA
 A  P  K  D  P  K  A  Q  P  R  F  D  Q  M  E  A  L  D  K  T  L  R  E  F  D  E

AAAGTCCGTGAAGTGCTCAAAAAATTGGGCTTTTCGGAAGAAGAAATCGAAAAATTCATAGAAGCTCTTAACAAGTAT
 K  V  R  E  V  L  K  K  L  G  F  S  E  E  E  I  E  K  F  I  E  A  L  N  K  Y

CTGGTGGACCAGTTCGTTAACCCGAAGCCGGAAGAGTTTGACATTGACAAGGTAGTAGAGGAAAAAGTCAAATTATTA
 L  V  D  Q  F  V  N  P  K  P  E  E  F  D  I  D  K  V  V  E  E  K  V  K  L  L

GAAGAATTT
 E  E  F
```

**Binder B8:**

```
ATGGGCTCGAAAAAAGAACTGGCGGAGAAATTCGTGGAACGTGCAGAAAAAGTGATCAAAGAGATTATTGAATCCGAA
 M  G  S  K  K  E  L  A  E  K  F  V  E  R  A  E  K  V  I  K  E  I  I  E  S  E

GAAGAAGTGTCCGAGAAAGATAAAGAGGTTCTGAAAATGTTGGTGAATGATATGAAACATTGGGCTGAACGAGCGGAG
 E  E  V  S  E  K  D  K  E  V  L  K  M  L  V  N  D  M  K  H  W  A  E  R  A  E

ACCCACAAAGAAATGGATGTGATTGACCGTTTTCAGTATGACATTCATATGCATGTTTGGGGTCTGGAGAACCCGAAA
 T  H  K  E  M  D  V  I  D  R  F  Q  Y  D  I  H  M  H  V  W  G  L  E  N  P  K

GAAGGCCTCGAAGACTATTTAATCCGTGAGATTCTTTCCGAATTT
 E  G  L  E  D  Y  L  I  R  E  I  L  S  E  F
```

**Binder B9:**

```
ATGGGCAAAGAAGAGGAGATTCAGAAAAAGATTGAAGAAATCATGGAGTTGATTAAAGAAGCTTATAAACTCGCACGC
 M  G  K  E  E  E  I  Q  K  K  I  E  E  I  M  E  L  I  K  E  A  Y  K  L  A  R

GAGTTTGCAGAAGCATCAGGCCTTGAAGAACTTAAACGCATGTTGTGGCATGTTGATTGGGAAATGCACTGGATCTTT
 E  F  A  E  A  S  G  L  E  E  L  K  R  M  L  W  H  V  D  W  E  M  H  W  I  F

CCGCAAAAACCGCCGACATTTGAAGACATAGAAAACGGTGTTATAGTGCTCGAGGTATTTGCCGAACGTGTGCGAGAA
 P  Q  K  P  P  T  F  E  D  I  E  N  G  V  I  V  L  E  V  F  A  E  R  V  R  E

CTGCTGAAAGATCCGAATCTGTCGGAAGAAGCGCGTAAAATCGCTGAAGAGTTTCTGGAGTTGTACGAAAAAATCAAA
 L  L  K  D  P  N  L  S  E  E  A  R  K  I  A  E  E  F  L  E  L  Y  E  K  I  K

AAAAAATATGAGGAACTGAAAAAGTTATTAGAGGAATTT
 K  K  Y  E  E  L  K  K  L  L  E  E  F
```

**Binder B10:**

```
ATGGGCAGTCTGGAAGAGGAAGTAGAAAAAGAAGTGGAAAAAGTGGCCGAAATTTTGAGCGATCCGAACCATACGCGT
 M  G  S  L  E  E  E  V  E  K  E  V  E  K  V  A  E  I  L  S  D  P  N  H  T  R

GAACAGTACAAGGAAGCTGTGTATAGCTTTTTCACCGCAATGGAACTGCAGAAGGAGGTCTATGAGAGCAAGGGAATT
 E  Q  Y  K  E  A  V  Y  S  F  F  T  A  M  E  L  Q  K  E  V  Y  E  S  K  G  I

AAGAAAACCCGTCAGGAAGTATTCGACGAAGTTGTAAAAAAAGTATGGGAAAAGGTTGAAGAGAAAGTTAAAGAACTG
 K  K  T  R  Q  E  V  F  D  E  V  V  K  K  V  W  E  K  V  E  E  K  V  K  E  L

AGCGAGAAAGATCCAGAAAAAGCCAAAAAAGCCGAGCACAAGGCACACATGTTCGATTACGATCTGGATTGGATTGCC
 S  E  K  D  P  E  K  A  K  K  A  E  H  K  A  H  M  F  D  Y  D  L  D  W  I  A

TGGGATAATCCAGACCTGACGCCGGAAGAGATCGTTGCGAAATACTTCCCAAAAGAATTT
 W  D  N  P  D  L  T  P  E  E  I  V  A  K  Y  F  P  K  E  F
```

**LUCOS adapter protein**

DNA sequence and translation of the LUCOS adapter protein containing B2 as competitor domain. For the different adapter protein, the competitor domain was replaced with B6. Color codes: 6xHis-tag highlighted in gray; LBit highlighted in blue; protein G dimer highlighted in yellow; amber codon (overridden for pBpA insertion) highlighted in green; semi-flexible linker highlighted in pink; *de novo* competitor domain B2 highlighted in teal; SBit ($K_D$ 2.5 μM) highlighted in cyan; Strep-Tag II sequence highlighted in red.

```
ATGGGCAGCAGCCATCATCATCATCATCACAGCAGCGGCCTGGTGCCGCGCGGCAGCCATATGGGAGGTTCGGTCTTC
 M  G  S  S  H  H  H  H  H  H  S  S  G  L  V  P  R  G  S  H  M  G  G  S  V  F

ACACTCGAAGATTTCGTTGGGGACTGGGAACAGACAGCCGCCTACAACCTGGACCAAGTCCTTGAACAGGGAGGTGTG
 T  L  E  D  F  V  G  D  W  E  Q  T  A  A  Y  N  L  D  Q  V  L  E  Q  G  G  V

TCCAGTTTGCTGCAGAATCTCGCCGTGTCCGTAACTCCGATCCAAAGGATTGTCCGGAGCGGTGAAAATGCCCTGAAG
 S  S  L  L  Q  N  L  A  V  S  V  T  P  I  Q  R  I  V  R  S  G  E  N  A  L  K

ATCGACATCCATGTCATCATCCCGTATGAAGGTCTGAGCGCCGACCAAATGGCCCAGATCGAAGAGGTGTTTAAGGTG
 I  D  I  H  V  I  I  P  Y  E  G  L  S  A  D  Q  M  A  Q  I  E  E  V  F  K  V

GTGTACCCTGTGGATGATCATCACTTTAAGGTGATCCTGCCCTATGGCACACTGGTAATCGACGGGGTTACGCCGAAC
 V  Y  P  V  D  D  H  H  F  K  V  I  L  P  Y  G  T  L  V  I  D  G  V  T  P  N

ATGCTGAACTATTTCGGACGGCCGTATGAAGGCATCGCCGTGTTCGACGGCAAAAAGATCACTGTAACAGGGACCCTG
 M  L  N  Y  F  G  R  P  Y  E  G  I  A  V  F  D  G  K  K  I  T  V  T  G  T  L

TGGAACGGCAACAAAATTATCGACGAGCGCCTGATCACCCCCGACGGCTCCATGCTGTTCCGAGTAACCATCAACAGC
 W  N  G  N  K  I  I  D  E  R  L  I  T  P  D  G  S  M  L  F  R  V  T  I  N  S

GGAGGCTCTGGAGGTTCTATGACATTCAAACTGATTATAAACGGCAAGACCCTGAAAGGGGAAATTACCATTGAGGCC
 G  G  S  G  G  S  M  T  F  K  L  I  I  N  G  K  T  L  K  G  E  I  T  I  E  A

GTGGACGCGTAGGAGGCGGAGAAGATCTTTAAACAATACGCAAATGACTATGGAATTGATGGGGAATGGACTTACGAC
 V  D  A  *  E  A  E  K  I  F  K  Q  Y  A  N  D  Y  G  I  D  G  E  W  T  Y  D

GACGCTACAAAAACCTTTACCGTGACGGAAGGTACCGGGGGGTCAGGTGGCTCAGGGGGAAGCGGTGGTTCCGGCGGT
 D  A  T  K  T  F  T  V  T  E  G  T  G  G  S  G  G  S  G  G  S  G  G  S  G  G

AGCGGAGGAAGCGGCGCTGAGGCTGCTGCCAAAGAGGCTGCAGCAAAAGAGGCAGCCGCGAAGGAGGCGGCGGCCAAA
 S  G  G  S  G  A  E  A  A  A  K  E  A  A  A  K  E  A  A  A  K  E  A  A  A  K

GAAGCGGCTGCCAAGGAGGCTGCCGCAAAAGCTGGTTCTGGGGGCAGCGGTGGTAGTGGCGGCAGTGGAGGCTCCGGA
 E  A  A  A  K  E  A  A  A  K  A  G  S  G  G  S  G  G  S  G  G  S  G  G  S  G

GGTAGCGGTGCCGAGGCAGCTGCAAAAGAAGCAGCAGCTAAAGAAGCTGCTGCAAAGGAGGCCGCAGCGAAGGAAGCT
 G  S  G  A  E  A  A  A  K  E  A  A  A  K  E  A  A  A  K  E  A  A  A  K  E  A

GCGGCTAAAGAGGCCGCTGCGAAGGCTGGCTCAGGCGGATCTGGCGGTTCTGGCGGAAGCGGCGGCTCTGGTGGATCT
 A  A  K  E  A  A  A  K  A  G  S  G  G  S  G  G  S  G  G  S  G  G  S  G  G  S

GGAGGCTGTACTAGTATGGGCAAAAAATCAGACGAAGAGGTCGGTCTCGAATTAGCTAAAGAGATTCTCAAGGAACTG
 G  G  C  T  S  M  G  K  K  S  D  E  E  V  G  L  E  L  A  K  E  I  L  K  E  L

ATTGAATCAATGGGGTTGAGCGAAGTTCCCAAAAAAGAGGAACTGGACCATTTCCTGCCGCACCTGGAATGGGACGCG
 I  E  S  M  G  L  S  E  V  P  K  K  E  E  L  D  H  F  L  P  H  L  E  W  D  A

GACTGGCAGATACAGGAACGTATTGTCGAATACTATAAAGAAAACGGAGAAGAACCGACGGAGGAACGCCTTGAAACA
 D  W  Q  I  Q  E  R  I  V  E  Y  Y  K  E  N  G  E  E  P  T  E  E  R  L  E  T
```

```
GCCCATAAAGCGGCGTGGTCTGTACTGCAGAAATTTATCGCGGAAGTCCGTGCCATGGCAGCAAAAGGCGATCCGTCG
 A  H  K  A  A  W  S  V  L  Q  K  F  I  A  E  V  R  A  M  A  A  K  G  D  P  S

AAAGAAGAAATTTTGGAGGTCATTGAATCCCTTGAAGCGGAATTTACAGGAGGTTCGGGTGGGTCGACCGGTAGCGTC
 K  E  E  I  L  E  V  I  E  S  L  E  A  E  F  T  G  G  S  G  G  S  T  G  S  V

ACCGGCTATCGGCTCTTTGAGAAAGAATCTGGAGGTTCTGGCGGATCTTGGTCCCATCCGCAGTTTGAAAAA
 T  G  Y  R  L  F  E  K  E  S  G  G  S  G  G  S  W  S  H  P  Q  F  E  K
```
